# A new set of composite, non-redundant electroencephalogram measures of non-rapid eye movement sleep based on the power law scaling of the Fourier spectrum

**DOI:** 10.1101/2020.04.09.035113

**Authors:** Róbert Bódizs, Orsolya Szalárdy, Csenge Horváth, Péter P. Ujma, Ferenc Gombos, Péter Simor, Adrián Pótári, Marcel Zeising, Axel Steiger, Martin Dresler

## Abstract

A novel method for deriving composite, non-redundant measures of non-rapid eye movement (NREM) sleep electroencephalogram (EEG) is developed on the basis of the power law scaling of the Fourier spectra. Measures derived are the spectral intercept, the slope (spectral exponent), as well as the maximal whitened spectral peak amplitude and frequency in the sleep spindle range. As a proof of concept, we apply these measures on a large sleep EEG dataset (N = 175; 81 females; age range: 17–60 years) with previously demonstrated effects of age, sex and intelligence. As predicted, aging is associated with decreased overall spectral slopes (increased exponents) and whitened spectral peak amplitudes in the spindle frequency range. In addition, age associates with decreased sleep spindle spectral peak frequencies in the frontal region. Women were characterized by higher spectral intercepts and higher spectral peak frequencies in the sleep spindle range. No sex differences in whitened spectral peak amplitudes of the sleep spindle range were found. Intelligence correlated positively with whitened spectral peak amplitudes of the spindle frequency range in women, but not in men. Last, age-related increases in spectral exponents did not differ in subjects with average and high intelligence. Our findings replicate and complete previous reports in the literature, indicating that the number of variables describing NREM sleep EEG can be effectively reduced in order to overcome redundancy and Type I statistical errors in future electrophysiological studies of sleep.

**Author summary:** Given the tight reciprocal relationship between sleep and wakefulness, the objective description of the complex neural activity patterns characterizing human sleep is of utmost importance in understanding the several facets of brain function, like sex differences, aging and cognitive abilities. Current approaches are either exclusively based on visual impressions expressed in graded levels of sleep depth (W, N1, N2, N3, REM), whereas computerized quantitative methods provide an almost infinite number of potential metrics, suffering from significant redundancy and arbitrariness. Our current approach relies on the assumptions that the spontaneous human brain activity as reflected by the scalp-derived electroencephalogram (EEG) are characterized by coloured noise-like properties. That is, the contribution of different frequencies to the power spectrum of the signal are best described by power law functions with negative exponents. In addition, we assume, that stages N2–N3 are further characterized by additional non-random (non-noise like, sinusoidal) activity patterns, which are emerging at specific frequencies, called sleep spindles (9–18 Hz). By relying on these assumptions we were able to effectively reduce 191 spectral measures to 4: (1) the spectral intercept reflecting the overall amplitude of the signal, (2) the spectral slope reflecting the constant ratio of low over high frequency power, (3) the frequency of the maximal sleep spindle activity and (4) the amplitude of the sleep spindle spectral peak. These 4 measures were efficient in characterizing known age-effects, sex-differences and cognitive correlates of sleep EEG. Future clinical and basic studies are supposed to be significantly empowered by the efficient data reduction provided by our approach.

## Introduction

The frequency characteristics of sleep-dependent neuronal oscillations as recorded by scalp electroencephalography (EEG) are increasingly recognized as potent markers of aging (Pótári et al., 2017; Ujma et al., 2019), health and disease (Kaskie and Ferrarelli, 2019), typical and atypical development and maturation (Campbell et al., 2012; Bódizs et al., 2012), as well as of neurocognitive features of high practical relevance (Bódizs et al., 2005; Ujma et al., 2017; Ujma, 2018). However, many of these studies are suffering from increased susceptibility to Type I error as a result of an inherently increased level of “researcher degrees of freedom”. That is, EEG data can be analysed in almost infinite different ways, by focusing on one or another specific electrophysiological phenomenon (Ujma, 2018; Ujma et al., 2020). Instead of focusing on multiple frequencies or phenomena, our aim is to provide an overall characterization of the broadband NREM sleep EEG. Our data-driven approach is based on the statistical properties of the signal, in order to assess the intercept and the slope, as well as the most prominent/important spectral peaks of the Fourier spectrum.

Evidence suggests the linear relationship between the logarithmic amplitude or power of EEG and the logarithm of frequency (Feinberg et al., 1984; Pereda et al., 1998). Such power law scaling is a general, state-independent feature of cortical EEG, suggesting that the Fourier spectrum can be reliably described by an approximation of the parameters of the following function:

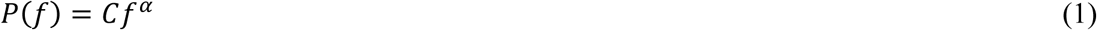

where *P* is power (*P* ≥ 0) as a function of frequency (0 ≤ *f* ≤ *f*_*Nyquist*_), C is the constant (or the intercept) expressing the overall, frequency-independent EEG amplitude (*C* > 0), whereas *α* is the spectral exponent indicating the decay rate (slope) of power as a function of frequency. Reported values for the spectral exponent are −4 < *α* < −1, with lower values indicating lower arousal/deeper sleep, but the values might depend on the EEG reference used, e.g. bipolar derivations result in higher α values as compared to referential ones, as well (Freemen et al., 2006; Lázár et al., 2020). That is, instead of providing 191 values for the power spectra of 0.5–48 Hz activity in bins of 0.25 Hz, we need just 2 (*C* and *α*). Most notably, if reliable, this function suggests that classical bandwise or binwise spectral analyses are not considering the frequency-determined nature of power values when applying statistical tests focusing on specific oscillatory phenomena.

However, there are further specific features of the EEG spectrum, known as spectral peaks, which are upward deflections in the decreasing power law trend described by function (1) above. These peaks reflect non-random oscillatory activities of specific frequencies, which might prevent the reliable estimation of α if they are not considered (Freeman and Zhai, 2009, Colombo et al., 2019). In order to deliberately describe the power spectrum by taking into account its prominent peaks, we suggest the inclusion of a peak power function in the formula as follows:

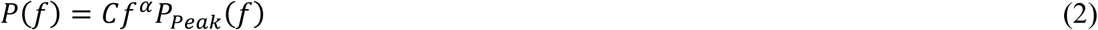

Peak power (*P*_*Peak*_) at frequency *f* equals 1 if there is no peak and is larger than 1 if there is a spectral peak at that frequency. Thus, the number of parameters is increased by considering spectral peaks, but is still lower than the values included in the original spectra, as putative “no peak regions” can be compressed in series of all ones. It has to be noted, that *P*_*Peak*_*(f)* is a whitened power measure, because it is independent from the spectral slope (*α*), which constitute the coloured part of the spectrum (Fig 1). In the following we only consider the maximal peaks, for which *P*_*Peak*_*(f)* ≤ *P*_*Peak*_*(f*_*maxPeak*_). No multiple peaks are analysed in this report.

**Fig 1.**
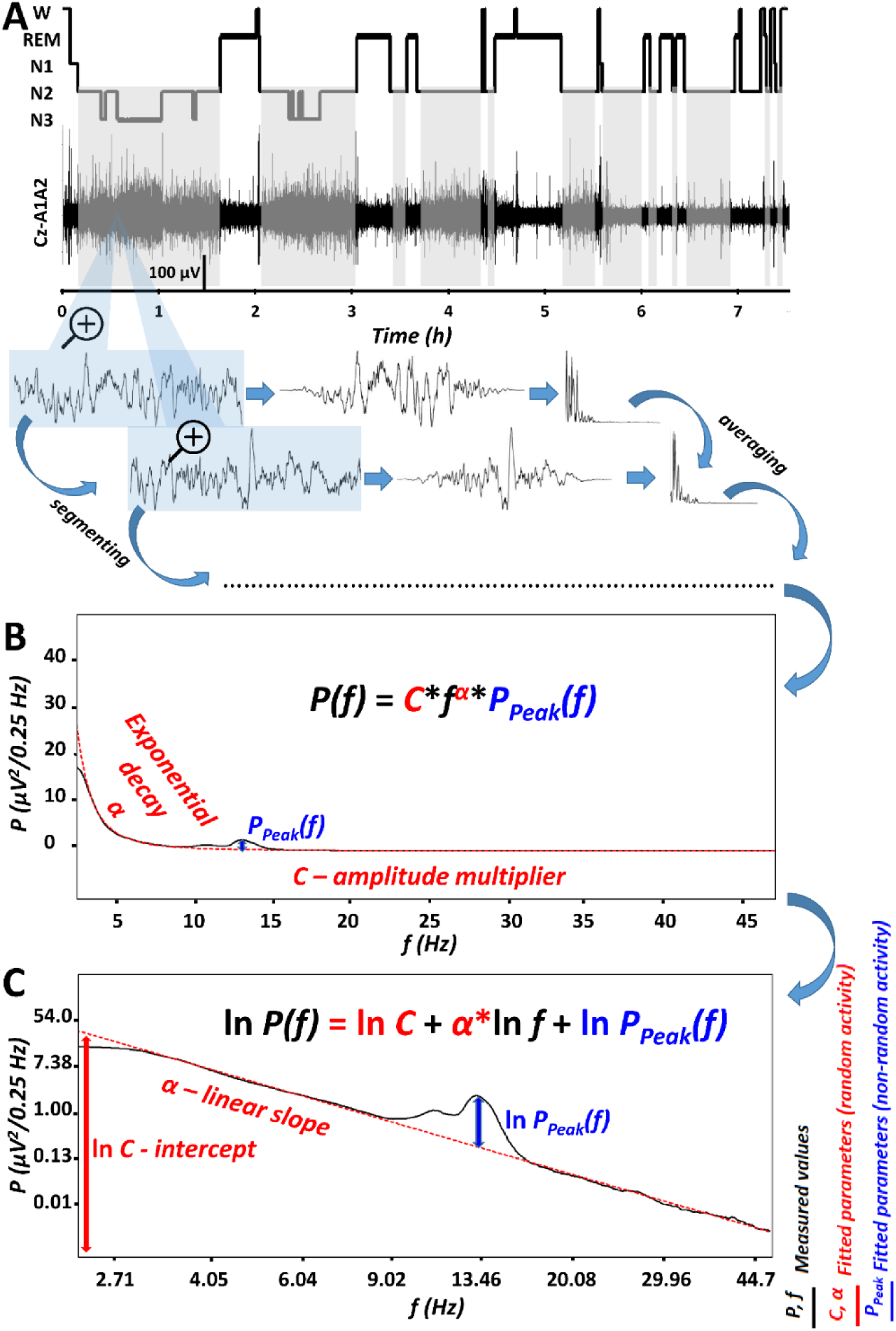
The parametrization of non-rapid eye movement (NREM) sleep electroencephalogram (EEG) spectra. A. Hypnogram and steps of the spectral EEG analyses as exemplified in a representative record of a young male volunteer. Grey shaded areas represent NREM sleep, which is analysed in the present report. Blue-shaded EEG segments are magnified 4 seconds long epochs, with 2 seconds overlap and modified with a Hanning window before power spectral analysis via mixed-radix Fast Fourier Transformation (FFT). B. Average spectral power (P) is characterized by a frequency (f)-dependent exponential decay (α), as well as by an overall, frequency-independent amplitude multiplier (C) and a peak power multiplier at critical frequencies [P_Peak_(f)]. C. The natural logarithm of spectral power (P) is a linear function of the natural logarithm of frequency (f), characterized by a linear slope α (which is equal with α in panel B) and an intercept (the latter being the natural logarithm of the amplitude multiplier, C in panel B). In addition, this linear function has to be summed with the natural logarithm of the peak power multiplier [P_Peak_(f), equal to the same frequency-dependent function in panel B]. Please note that “no peak regions” can be compressed in series of all ones, resulting in reduced number of variables as compared to the bins in the original spectra.

As a proof of concept, we apply these measures on a large sleep EEG dataset with previously demonstrated effects of individual differences. Specifically, we translate some core findings regarding age, sex, and general intelligence-related effects in NREM sleep EEG into specific hypotheses in terms of *C, α*, and two indexes of spindle-range *P*_*Peak*_*(f)*: whitened spectral peak amplitude *P*_*Peak*_*(f*_*maxPeak*_) and spectral peak frequency (*f*_*maxPeak*_).

Age was reported to correlate negatively with NREM sleep EEG slow wave activity, but positively with high frequency activity in healthy adult subjects (Carrier et al., 2001). An early study based on period amplitude analysis reported that NREM sleep EEG log amplitude is a linear function of log frequency and that the slope of this linear decay is steeper in young as compared to older adults (Feinberg et al., 1984). Thus, we hypothesize (H1) that the slope of the Fast Fourier Transformation (FFT)-based spectrum of NREM sleep EEG (*α*) is age-dependently increasing (less steep decreasing trends are indexed by higher exponents *α*). In addition, aging was shown to be associated with decreased sleep spindle activity (Nicolas et al., 2001; Purcell et al., 2017), thus we hypothesize (H2) a negative correlation between age and *P*_*Peak*_*(f*_*maxPeak*_) values characterizing maximal spindle frequency spectral peaks (9 Hz < *f* < 18 Hz). In addition to decreased spindle activity, the increase in intra-spindle oscillatory frequency (Hz) was shown to be a characteristic feature of aging according to some (Principe and Smith, 1982; Nicolas et al., 2001), but not all (Purcell et al., 2017) reports. As a consequence, we hypothesize (H3) that the maxima of the *P*_*Peak*_*(f)* function for 9 Hz < *f* < 18 Hz (broad spindle range) emerge at higher *f*_*maxPeak*_ values in aged, as compared to young subjects.

Reported sex differences in NREM and REM sleep EEG indicate higher spectral power in several frequency bands in women, as compared to men (Dijk et al., 1989; Carrier et al., 2001). Such broad band and state-independent differences suggest a general tendency for a higher EEG amplitude in women, due to non-neuronal factors, like skull thickness and bone mineral density (Dijk et al., 1989; Looker et al., 2009). As a consequence, we hypothesize (H4) that women are characterized by higher spectral intercepts, than men (*C*_♀_ > *C*_♂_). As a consequence of this point we will reanalyze some of the reported sex differences in sleep spindle density/power, indicating increased sleep spindling in women as compared to men (Dijk et al., 1989; Carrier et al., 2001; Crowley, 2002; Huupponen et al., 2002), by relying on whitened spectral peak amplitudes of the spindle range (*P*_*Peak*_*(f*_*maxPeak*_)).

Another sex difference was reported in terms of sleep spindle frequency, that is women were shown to be characterized by higher oscillatory frequencies as compared to men (Ujma et al., 2014). We hypothesize (H5) that 9–18 Hz *P*_*Peak*_*(f)* maxima occurs at higher *f* values in women as compared to men (*f*_*maxPeak*♀_ > *f*_*maxPeak*♂_).

Intelligence was shown to correlate positively with NREM sleep EEG sleep spindle activity (Bódizs et al., 2005). Although, a recent metaanalysis casts doubt on the sexual dimorphism of this relationship (Ujma, 2018), the dataset we analyse in our current report is characterized by a clear difference among women and men: women were characterized by positive correlation between sleep spindle amplitude/power and IQ, whereas null correlations were reported for men (Ujma et al., 2014; Ujma et al., 2017). As our current analyses are based on the same dataset, we hypothesize (H6) that *P*_*Peak*_*(f*_*maxPeak*_) values of the sleep spindle range (9–18 Hz) correlate positively with IQ in women, but not in men. Intelligence was also reported to modulate the relationship between the decrease in NREM sleep EEG slow activity associated with aging: participants showing average IQ (AIQ) scores were characterized by significant negative correlations regarding age vs. slow wave activity, whereas no such correlations were found in individuals with high IQ (HIQ) (Pótári et al., 2017). As the original report provided overwhelming evidence for an age vs relative delta power correlation as being modulated by IQ range, whereas weaker evidence was found for absolute power (Pótári et al., 2017), we do not know if this finding reflects the age-dependency of slow wave activity per se, or the combined age-dependency of slow wave activity and slow/high activity ratio. The former scenario would fit with a null effect for IQ-modulation of age vs spectral slope correlation, whereas the latter would lead to an IQ-dependence of age vs spectral slope relationship (H7).

## Results

### Goodness of fit: Is the logarithm of spectral power a linear function of the logarithm of frequency?

Linears were fitted to the equidistant log-log plots of the EEG power spectra in the 2– 48 Hz range, excluding the 6–18 Hz range, the latter known to be characterized by spectral peaks in NREM sleep (Fig. 1C, see details in section Methods). The sample mean of fitted slopes 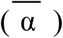 varied between −2.73 (SD = 0.22) and −2.33 (SD = 0.22) for the frontocentral (Fz) and left posterior temporal (T5) region, respectively. In turn, the sample mean of the intercepts 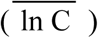 varied between 3.74 (SD = 0.73) and 5.76 (SD = 0.69) for derivations T5 and Fz, respectively. Goodness of fit (*R*^*2*^) of the linear model of the equidistant 2–6 Hz and 18–48 Hz spectral data varied in the range of 0.8955–0.9997 across subjects and EEG derivations. The square of the Fisher Z-transformed, averaged and back-transformed Pearson correlations between the fitted linear and the spectral data is 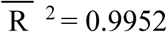 (SD = 0.1578).

#### H1: Age-associated increase in the spectral exponent (decrease in spectral slope) of NREM sleep EEG

Spearman rank correlations (ρ) indicated a significant positive association between age (years) and NREM sleep EEG spectral exponents (*α*) at all derivations (Table 1a). The Rüger’s area including all derivations proved to be significant at both of the new critical probability (*p*) levels (0.025 and 0.017). Thus, based on the Descriptive Data Analysis (DDA, see details in section Methods) procedure (Abt, 1987; Abt, 1990), this area can be considered as a significant one (see also Fig 2A).

**Table 1.**
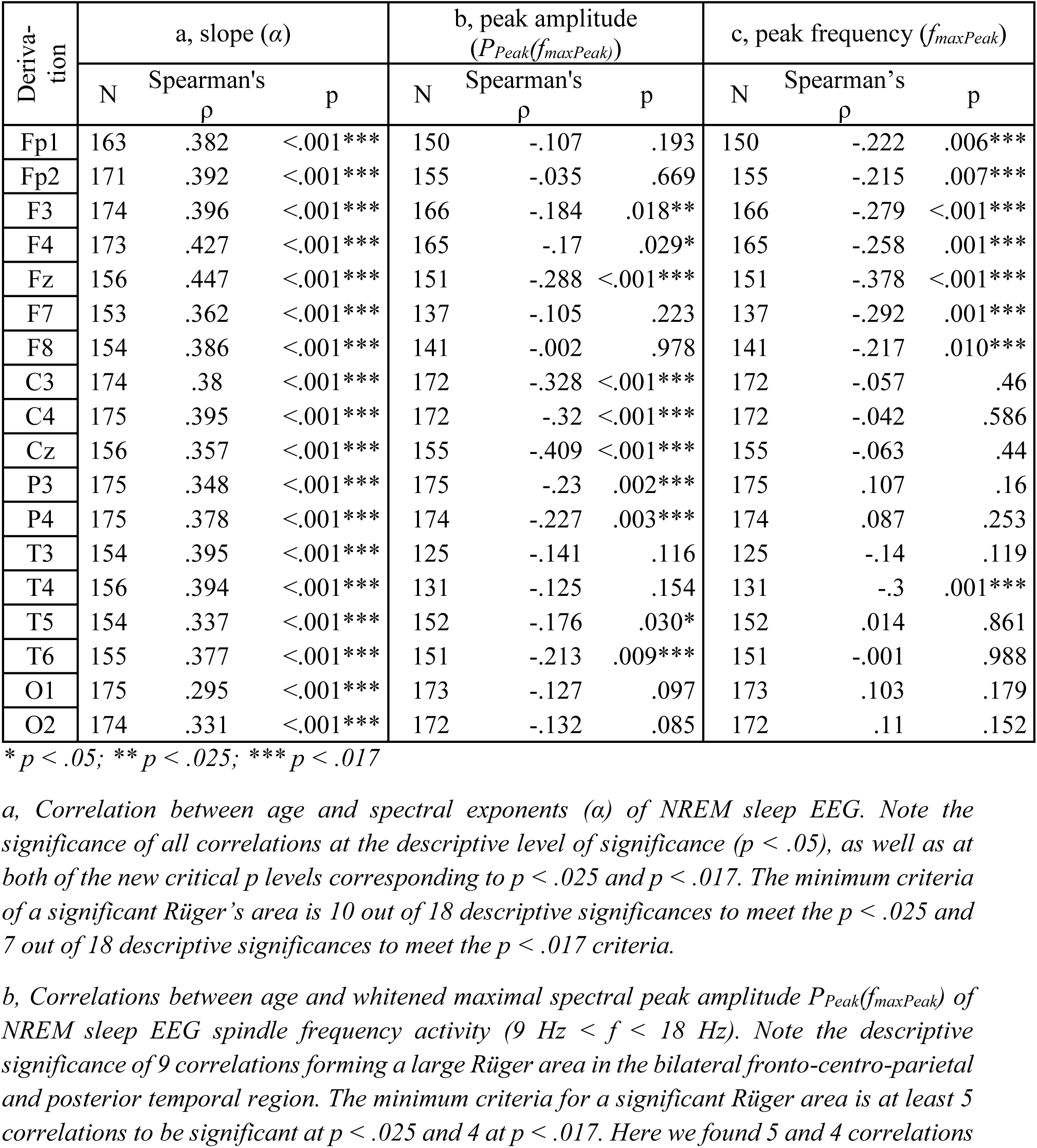
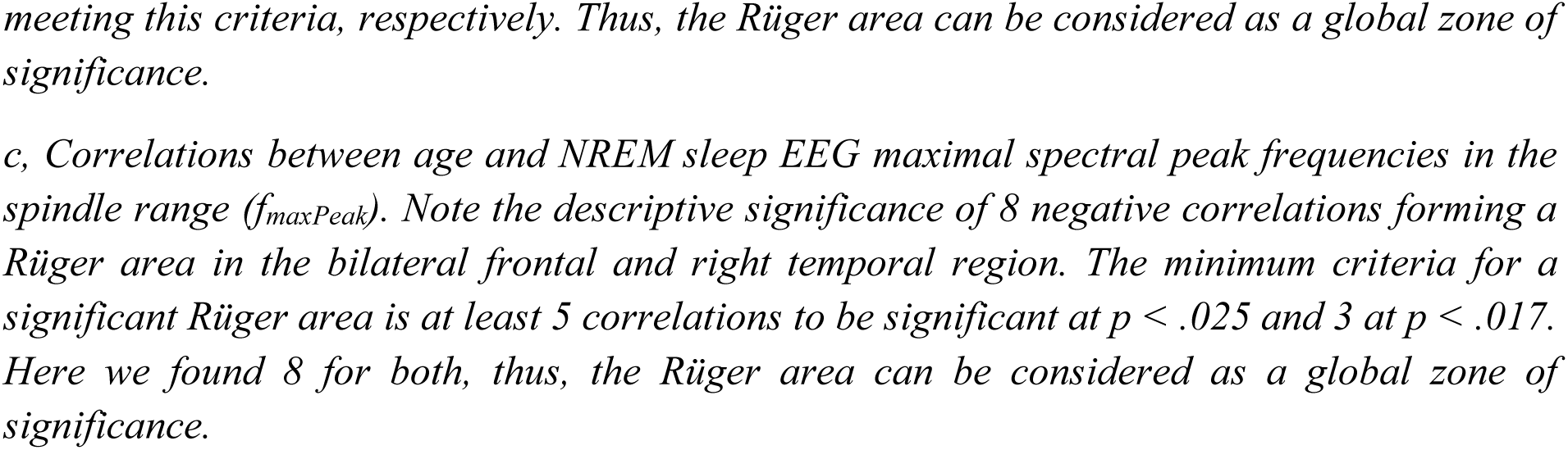
Spearman rank correlations between age and spectral exponents, as well as spectral peak amplitudes and frequencies in the sleep spindle range of NREM sleep EEG

| Derivation | a, slope ($\alpha$) | | | b, peak amplitude ($P_{Peak}(f_{maxPeak})$) | | | c, peak frequency ($f_{maxPeak}$) | | |
| --- | --- | --- | --- | --- | --- | --- | --- | --- | --- |
| | N | Spearman's $\rho$ | p | N | Spearman's $\rho$ | p | N | Spearman's $\rho$ | p |
| Fp1 | 163 | .382 | <.001*** | 150 | -.107 | .193 | 150 | -.222 | .006*** |
| Fp2 | 171 | .392 | <.001*** | 155 | -.035 | .669 | 155 | -.215 | .007*** |
| F3 | 174 | .396 | <.001*** | 166 | -.184 | .018** | 166 | -.279 | <.001*** |
| F4 | 173 | .427 | <.001*** | 165 | -.17 | .029* | 165 | -.258 | .001*** |
| Fz | 156 | .447 | <.001*** | 151 | -.288 | <.001*** | 151 | -.378 | <.001*** |
| F7 | 153 | .362 | <.001*** | 137 | -.105 | .223 | 137 | -.292 | .001*** |
| F8 | 154 | .386 | <.001*** | 141 | -.002 | .978 | 141 | -.217 | .010*** |
| C3 | 174 | .38 | <.001*** | 172 | -.328 | <.001*** | 172 | -.057 | .46 |
| C4 | 175 | .395 | <.001*** | 172 | -.32 | <.001*** | 172 | -.042 | .586 |
| Cz | 156 | .357 | <.001*** | 155 | -.409 | <.001*** | 155 | -.063 | .44 |
| P3 | 175 | .348 | <.001*** | 175 | -.23 | .002*** | 175 | .107 | .16 |
| P4 | 175 | .378 | <.001*** | 174 | -.227 | .003*** | 174 | .087 | .253 |
| T3 | 154 | .395 | <.001*** | 125 | -.141 | .116 | 125 | -.14 | .119 |
| T4 | 156 | .394 | <.001*** | 131 | -.125 | .154 | 131 | -.3 | .001*** |
| T5 | 154 | .337 | <.001*** | 152 | -.176 | .030* | 152 | .014 | .861 |
| T6 | 155 | .377 | <.001*** | 151 | -.213 | .009*** | 151 | -.001 | .988 |
| O1 | 175 | .295 | <.001*** | 173 | -.127 | .097 | 173 | .103 | .179 |
| O2 | 174 | .331 | <.001*** | 172 | -.132 | .085 | 172 | .11 | .152 |

\* $p < .05$ ; \*\* $p < .025$ ; \*\*\* $p < .017$
*a, Correlation between age and spectral exponents ( $\alpha$ ) of NREM sleep EEG. Note the significance of all correlations at the descriptive level of significance ( $p < .05$ ), as well as at both of the new critical $p$ levels corresponding to $p < .025$ and $p < .017$ . The minimum criteria of a significant Rger's area is 10 out of 18 descriptive significances to meet the $p < .025$ and 7 out of 18 descriptive significances to meet the $p < .017$ criteria.*
*b, Correlations between age and whitened maximal spectral peak amplitude $P_{Peak}(f_{maxPeak})$ of NREM sleep EEG spindle frequency activity ( $9 \text{ Hz} < f < 18 \text{ Hz}$ ). Note the descriptive significance of 9 correlations forming a large Rger area in the bilateral fronto-centro-parietal and posterior temporal region. The minimum criteria for a significant Rger area is at least 5 correlations to be significant at $p < .025$ and 4 at $p < .017$ . Here we found 5 and 4 correlations*
meeting this criteria, respectively. Thus, the Rger area can be considered as a global zone of significance.
c, Correlations between age and NREM sleep EEG maximal spectral peak frequencies in the spindle range ( $f_{\text{maxPeak}}$ ). Note the descriptive significance of 8 negative correlations forming a Rger area in the bilateral frontal and right temporal region. The minimum criteria for a significant Rger area is at least 5 correlations to be significant at $p < .025$ and 3 at $p < .017$ . Here we found 8 for both, thus, the Rger area can be considered as a global zone of significance.

**Fig 2.**
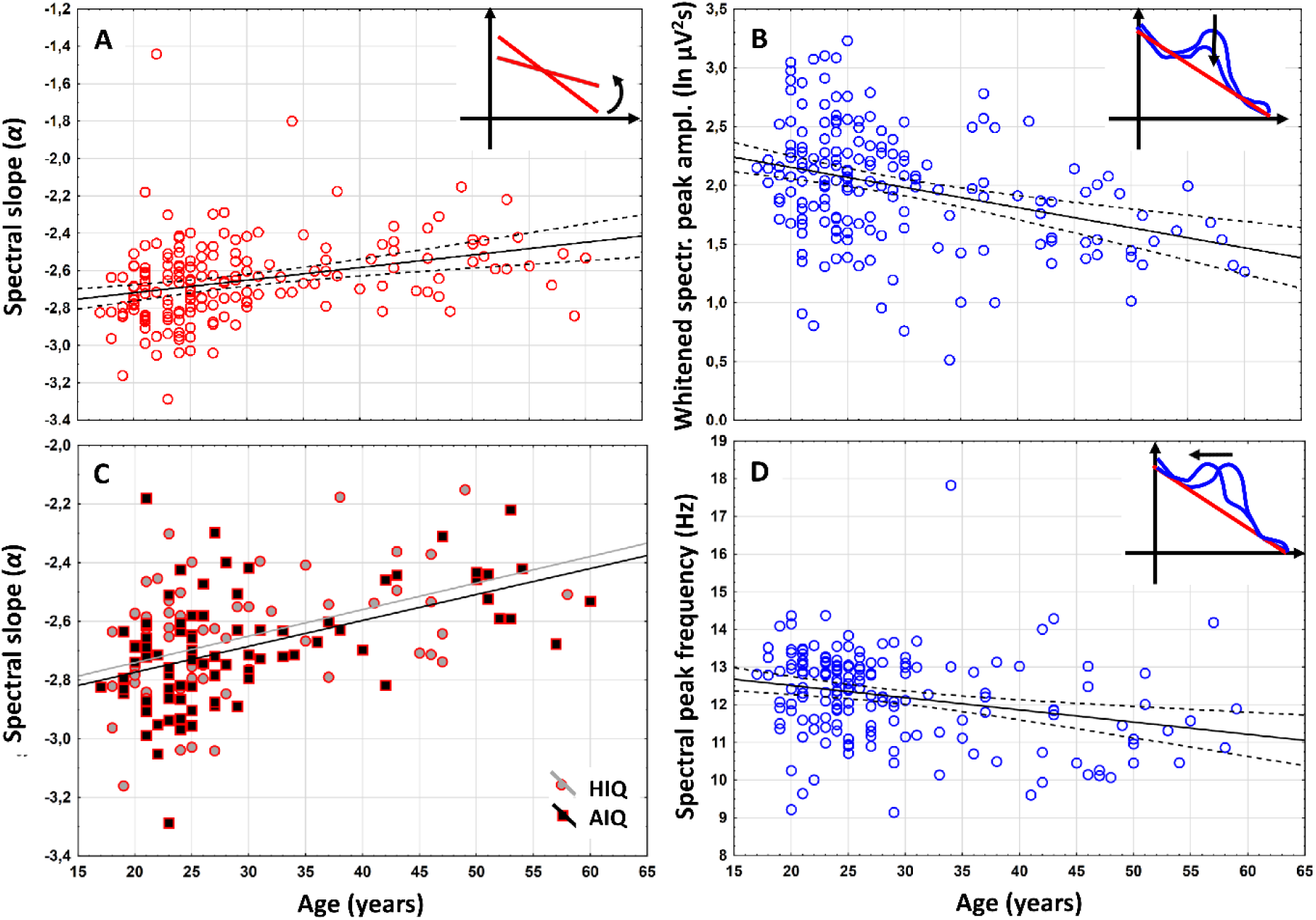
Representative scatterplots of the correlations between age and measures of the NREM sleep EEG spectra at derivation F3. A. Correlation of age with the spectral exponent (α) indicating a decrease in the spectral slope in the aged subjects. B. Correlations of age with the whitened maximal spectral peak amplitude in the sleep spindle frequency range (P_Peak_(f_maxPeak_). Note the decrease in whitened spectral peak amplitude in the aged. C. Correlation of age with the NREM sleep EEG spectral exponent (α) as categorized by intelligence (HIQ – High Intelligence Quotient, AIQ – Average Intelligence Quotient). Note the lack of an IQ effect. D. Correlation of age with NREM sleep EEG maximal spectral peak frequency (f_maxPeak_) in the spindle range. Note the age-dependent decline in frequency. Color codes are consistent with Fig 1: red –spectral slopes, blue – spectral peaks.

#### H2: Age-dependent decrease in whitened spectral peak amplitude of NREM sleep EEG sleep spindle frequencies

Based on Spearman’s rank correlations (ρ), maximal whitened spectral peak amplitudes of NREM sleep EEG spindle frequencies (*P*_*Peak*_*(f*_*maxPeak*_)) and age correlated negatively at 10 derivations covering the frontocentral, parietal and posterior temporal areas (F3, F4, Fz, C3, Cz, C4, T5, T6, P3, and P4). Among these 10 derivations defining the Rüger area based on descriptive significances, 8 were significant at *p* < .025 and 7 at *p* < .017 (Table 1b). That is the Rüger area indicates a negative correlation between age and whitened sleep spindle spectral peak amplitude (see a representative example in Fig 2B).

#### H3: Age-related decrease but not increase in spectral peak frequency of NREM sleep EEG spindle range activity was found

Based on Spearman’s rank correlation (ρ) maximal sleep spindle spectral peak emerge at lower *f*_*maxPeak*_ values in the frontal region of aged, as compared to young subjects. This finding evidently contrasts our prediction. Peak frequency and age correlated negatively at 8 derivations covering the frontal and the right temporal areas (Fp1, Fp2, F3, F4, Fz, F7, F8 and T4). This Rüger’s area was significant, as all correlations conformed to both of the new critical probabilities (Table 1c, Fig 3D).

**Fig 3.**
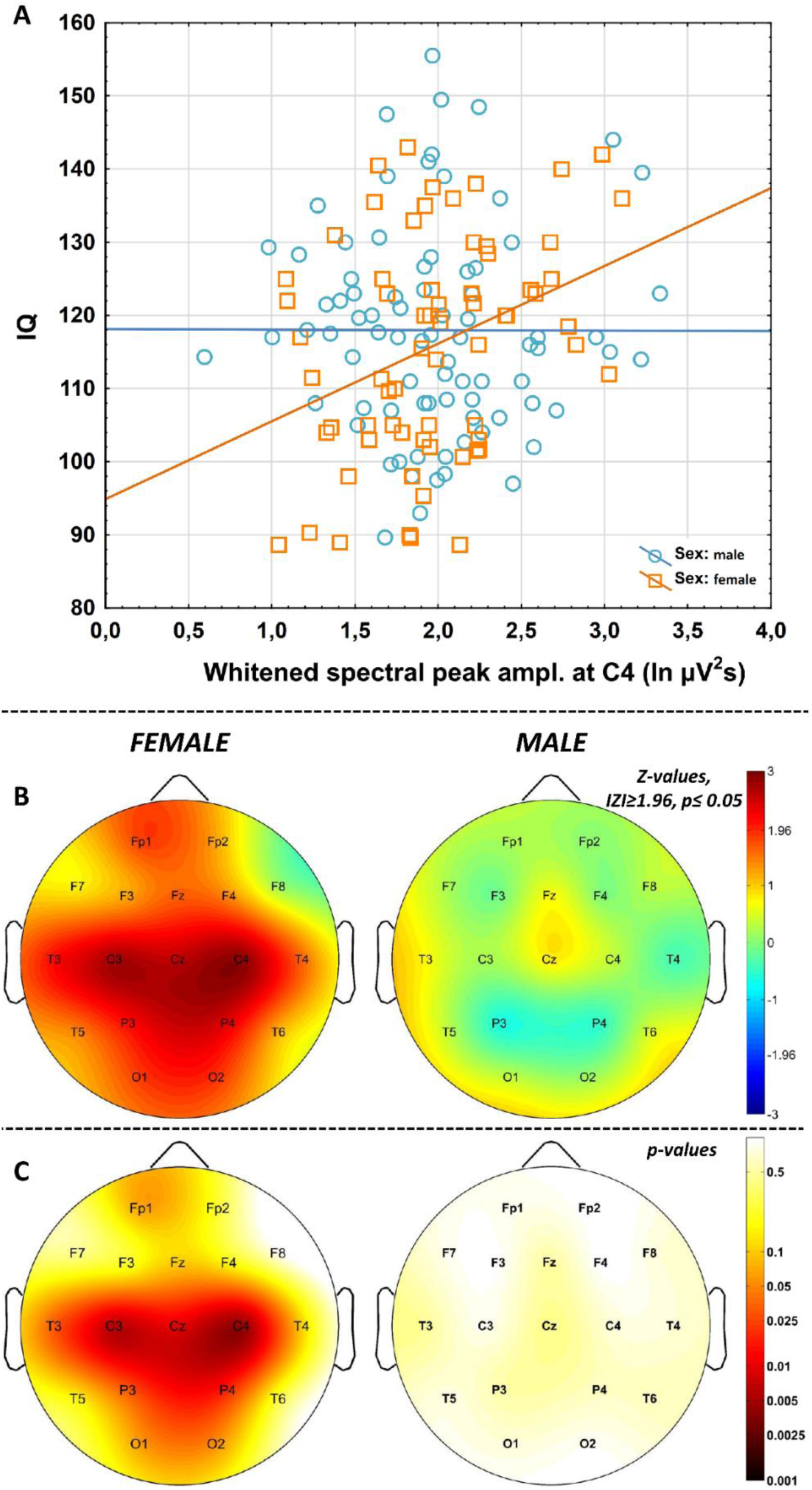
Correlations between NREM sleep EEG spindle frequency whitened spectral peak amplitudes and IQ in females and males. A. Categorized scatterplot representing the correlation between whitened spectral peak amplitude of the NREM sleep EEG spindle frequency range (recording site: F4) and IQ in women and men. B. Pearson r-values were transformed to Z-values and represented on topographical maps. C. Significance probability maps of the correlations presented in B.

#### H4: NREM sleep EEG spectral intercepts, but not whitened spindle spectral peak amplitudes are higher in women as compared to men

Mann-Whitney U test revealed that women are characterized by significantly higher spectral intercepts (the natural logarithm of *C* values in formula (1) and (2)) compared to men at all derivations. After correction for multiple testing the Rüger-area remained significant (Table 2). As predicted women and men did not differ in NREM sleep EEG maximal spectral peak amplitudes of the spindle range (*P*_*Peak*_*(f*_*maxPeak*_)) at any of the derivations (Table 2b).

**Table 2.** Women vs men differences in NREM sleep EEG spectral intercepts and whitened peak amplitudes in the spindle range

| | a, intercept (ln C) | | | | | | b, peak amplitude ($P_{Peak}(f_{maxPeak})$) | | | | | |
| --- | --- | --- | --- | --- | --- | --- | --- | --- | --- | --- | --- | --- |
| | U | p | N<sub>♀</sub> | Md<sub>♀</sub> | N<sub>♂</sub> | Md<sub>♂</sub> | t | p | N<sub>♀</sub> | $\bar{x}_{♀}$ | N<sub>♂</sub> | $\bar{x}_{♂}$ |
| Fp1 | 2455 | .004*** | 77 | 5.11 | 86 | 4.84 | -.098 | .924 | 67 | 1.370 | 83 | 1.379 |
| Fp2 | 2682 | .003*** | 81 | 5.14 | 90 | 4.87 | .515 | .619 | 68 | 1.404 | 87 | 1.361 |
| F3 | 2658 | .001*** | 81 | 5.53 | 93 | 5.22 | .420 | .675 | 75 | 1.731 | 91 | 1.699 |
| F4 | 2503 | <.001*** | 81 | 5.58 | 92 | 5.28 | 1.003 | .317 | 76 | 1.760 | 89 | 1.684 |
| Fz | 1997 | .001*** | 69 | 6.04 | 87 | 5.61 | .836 | .405 | 66 | 1.804 | 85 | 1.736 |
| F7 | 2295 | .031* | 67 | 4.58 | 86 | 4.28 | 1.474 | .162 | 59 | 1.470 | 78 | 1.336 |
| F8 | 2382 | .046* | 69 | 4.56 | 85 | 4.44 | .710 | .493 | 63 | 1.388 | 78 | 1.327 |
| C3 | 2345 | <.001*** | 81 | 5.36 | 93 | 4.98 | -.059 | .953 | 80 | 1.989 | 92 | 1.994 |
| C4 | 2483 | <.001*** | 81 | 5.41 | 94 | 5.04 | .225 | .822 | 79 | 1.983 | 93 | 1.966 |
| Cz | 1869 | <.001*** | 69 | 5.98 | 87 | 5.58 | .434 | .665 | 68 | 2.145 | 87 | 2.108 |
| P3 | 2467 | <.001*** | 81 | 4.97 | 94 | 4.69 | -.397 | .692 | 81 | 2.241 | 94 | 2.274 |
| P4 | 2423 | <.001*** | 81 | 5.04 | 94 | 4.56 | -.704 | .483 | 81 | 2.144 | 93 | 2.202 |
| T3 | 2270 | .019** | 67 | 3.99 | 87 | 3.72 | 1.028 | .306 | 55 | 1.254 | 70 | 1.169 |
| T4 | 2428 | .041* | 69 | 4.02 | 87 | 3.83 | .984 | .327 | 57 | 1.197 | 74 | 1.119 |
| T5 | 2172 | .006*** | 68 | 3.92 | 86 | 3.67 | -1.095 | .275 | 68 | 1.545 | 84 | 1.631 |
| T6 | 2021 | .001*** | 68 | 3.91 | 87 | 3.65 | -.460 | .646 | 66 | 1.475 | 85 | 1.509 |
| O1 | 2182 | <.001*** | 81 | 4.51 | 94 | 3.97 | -1.072 | .285 | 80 | 1.692 | 93 | 1.783 |
| O2 | 2102 | <.001*** | 81 | 4.53 | 93 | 4.04 | -1.938 | .054 | 80 | 1.592 | 92 | 1.748 |

a. Mann-Whitney U tests indicate higher spectral intercepts (ln C values) in the female (♀) as compared to the male (♂) subgroup. Descriptive significance was observed at all (18) recording locations. Minimum criteria of a significant Rger area requires at least 10 of these p values to be lower than .025 and 7 of them to be less than .017. Observed values are 15 and 14, respectively. Thus, the Rger area characterizing the sex differences in NREM sleep EEG spectral intercepts is significant. Md – median; \* $p < .05$ ; \*\* $p < .025$ ; \*\*\* $p < .017$ .
b. Whitened maximal spectral peak amplitudes in the sleep spindle range of NREM sleep EEG ( $P_{Peak}(f_{maxPeak})$ ) do not significantly differ between females and males.

#### H5: Women are characterized by higher NREM sleep EEG spectral peak frequencies in the spindle range

According to Mann-Whitney U Tests, women were characterized by significantly higher *f*_*maxPeak*_ values as compared to men, except derivations T3 and T4. The area remained significant after the correction for multiple testing (Table 3).

**Table 3.**
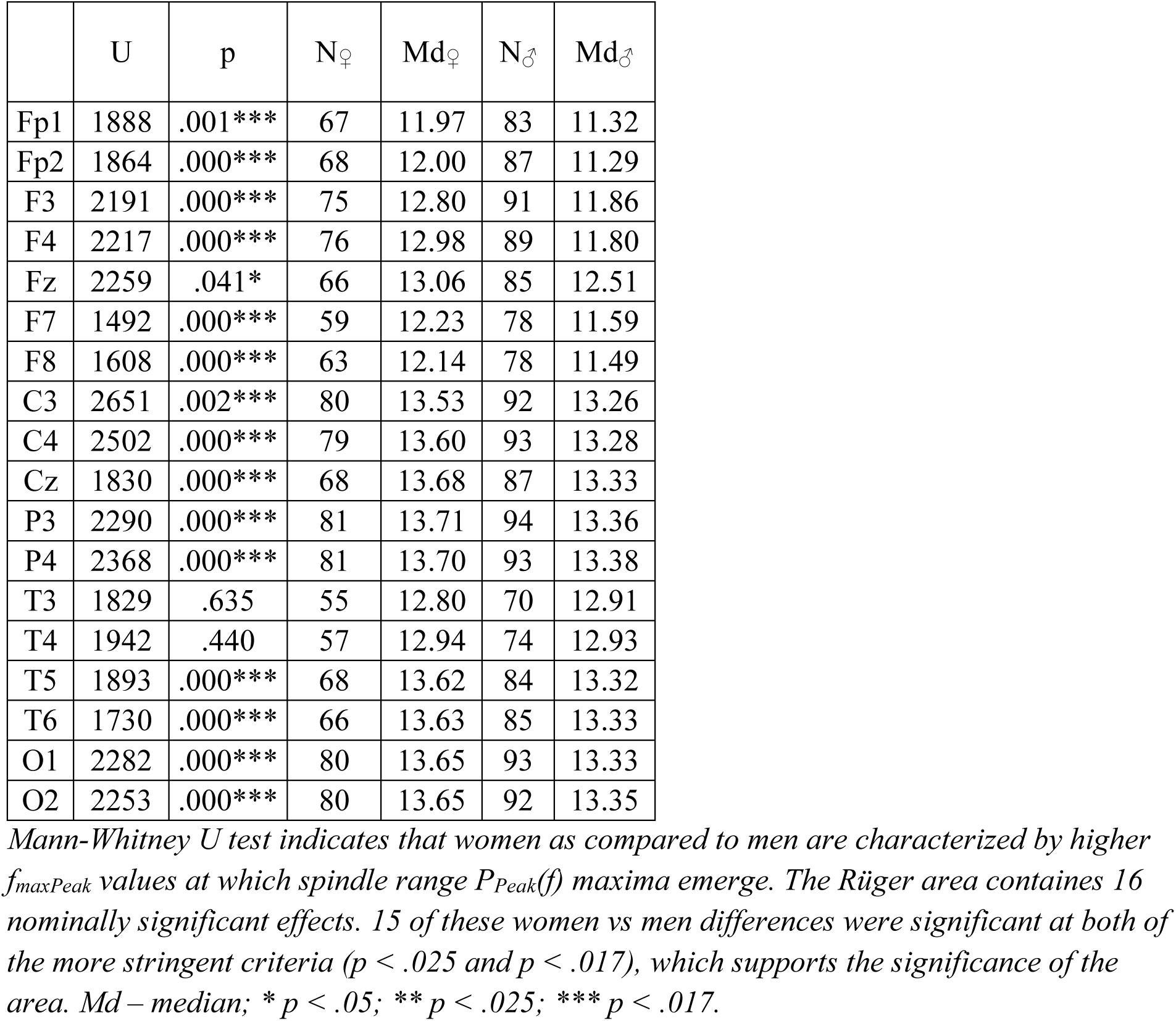
Women vs men differences in NREM sleep EEG spindle spectral peak frequencies

#### H6: IQ correlates positively with NREM sleep EEG spindle range whitened spectral peak amplitude in women

Pearson correlations revealed significant associations of whitened maximal spectral peak amplitudes (*P*_*Peak*_*(f*_*maxPeak*_)) pertaining to NREM sleep EEG spindle activity with IQ at derivations C3 (N = 67, r = .33, p = .007), C4 (N = 66, r = .34, p = .005), Cz (N = 55, r = .34, p = .010), P3 (N = 68, r = .26, p = .031), P4 (N = 68, r = .28, p = .020), and T3 (N = 45, r = .32, p = .031) in women (Table 4; Fig 3). The Rüger area at this centroparietal-left temporal region remained significant after the control for multiple testing (4/6 correlations are significant at .05/2 and 3/6 correlations at .05/3). No significant correlations of whitened spectral peak amplitude and IQ were found in men.

**Table 4.**
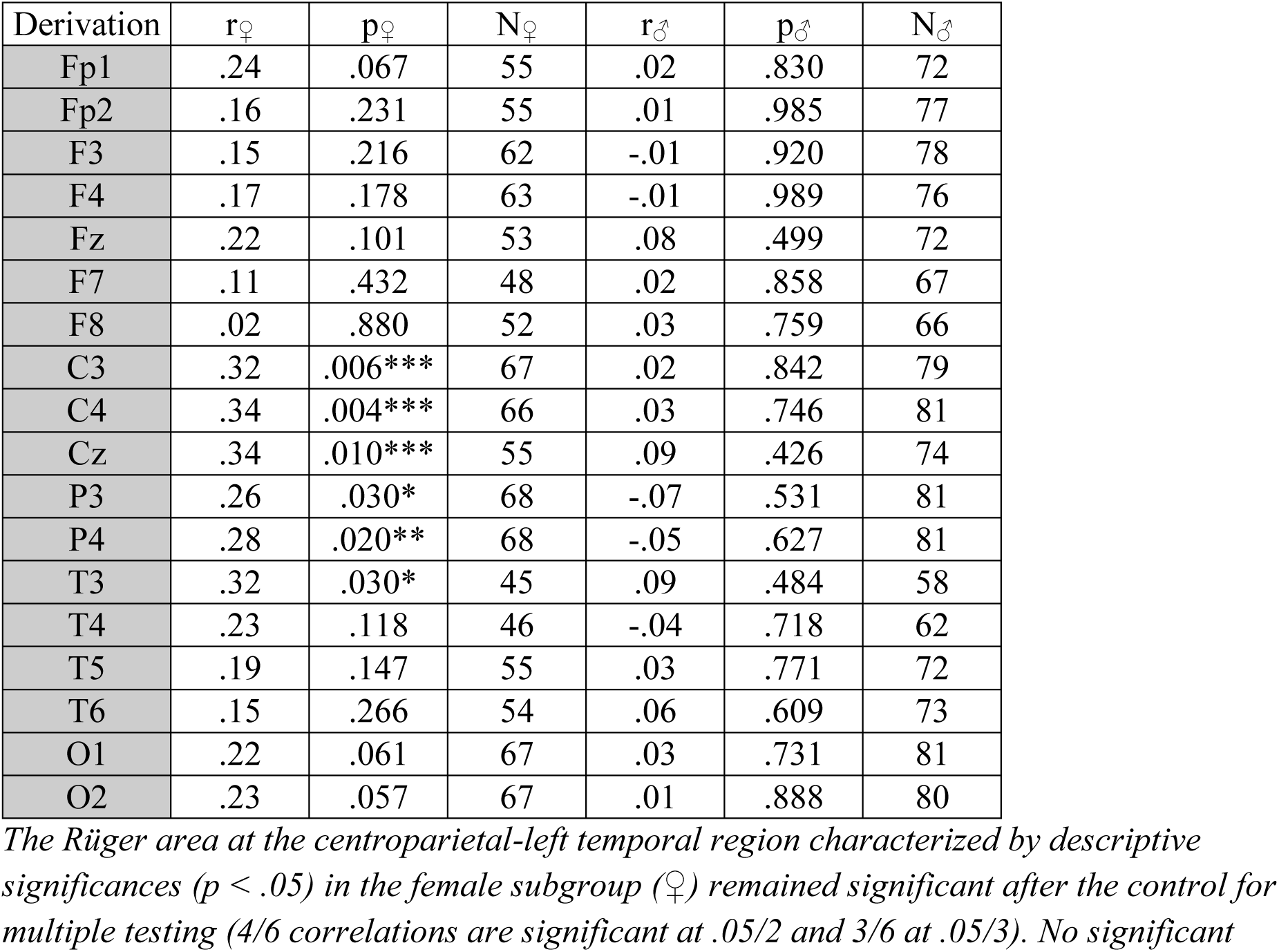
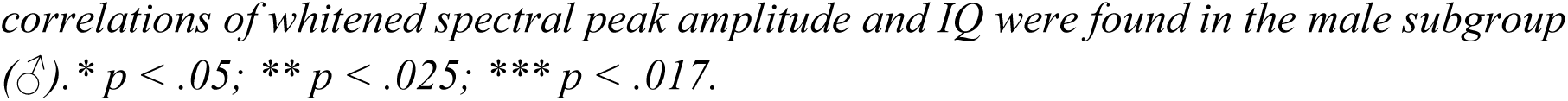
Correlation of whitened spectral peak amplitudes with IQ in women and men

| Derivation | $r_{\text{♀}}$ | $p_{\text{♀}}$ | $N_{\text{♀}}$ | $r_{\text{♂}}$ | $p_{\text{♂}}$ | $N_{\text{♂}}$ |
| --- | --- | --- | --- | --- | --- | --- |
| Fp1 | .24 | .067 | 55 | .02 | .830 | 72 |
| Fp2 | .16 | .231 | 55 | .01 | .985 | 77 |
| F3 | .15 | .216 | 62 | -.01 | .920 | 78 |
| F4 | .17 | .178 | 63 | -.01 | .989 | 76 |
| Fz | .22 | .101 | 53 | .08 | .499 | 72 |
| F7 | .11 | .432 | 48 | .02 | .858 | 67 |
| F8 | .02 | .880 | 52 | .03 | .759 | 66 |
| C3 | .32 | .006*** | 67 | .02 | .842 | 79 |
| C4 | .34 | .004*** | 66 | .03 | .746 | 81 |
| Cz | .34 | .010*** | 55 | .09 | .426 | 74 |
| P3 | .26 | .030* | 68 | -.07 | .531 | 81 |
| P4 | .28 | .020** | 68 | -.05 | .627 | 81 |
| T3 | .32 | .030* | 45 | .09 | .484 | 58 |
| T4 | .23 | .118 | 46 | -.04 | .718 | 62 |
| T5 | .19 | .147 | 55 | .03 | .771 | 72 |
| T6 | .15 | .266 | 54 | .06 | .609 | 73 |
| O1 | .22 | .061 | 67 | .03 | .731 | 81 |
| O2 | .23 | .057 | 67 | .01 | .888 | 80 |

*The Rger area at the centroparietal-left temporal region characterized by descriptive significances ( $p < .05$ ) in the female subgroup (♀) remained significant after the control for multiple testing (4/6 correlations are significant at .05/2 and 3/6 at .05/3). No significant*
correlations of whitened spectral peak amplitude and IQ were found in the male subgroup ( $\sigma$ ). \* $p < .05$ ; \*\* $p < .025$ ; \*\*\* $p < .017$ .

#### H7: Do age-related declines in NREM sleep EEG spectral slopes differ among subjects with average and high IQ?

As already presented in the former subheadings (H1) an age-associated increase in spectral exponents (less steep spectral slopes) characterizes the NREM sleep EEG of adult volunteers. This effect was separately assessed in subjects with average and high IQ, and results were compared. Age and slopes of the NREM sleep EEG spectra (*α*) were significantly associated in both subgroups (AIQ and HIQ). We found no significant difference between these correlations, however (Table 5). That is, age-associated decreases in the steepness of the slopes of the NREM sleep EEG spectra are independent of the subjects’ IQ.

**Table 5.**
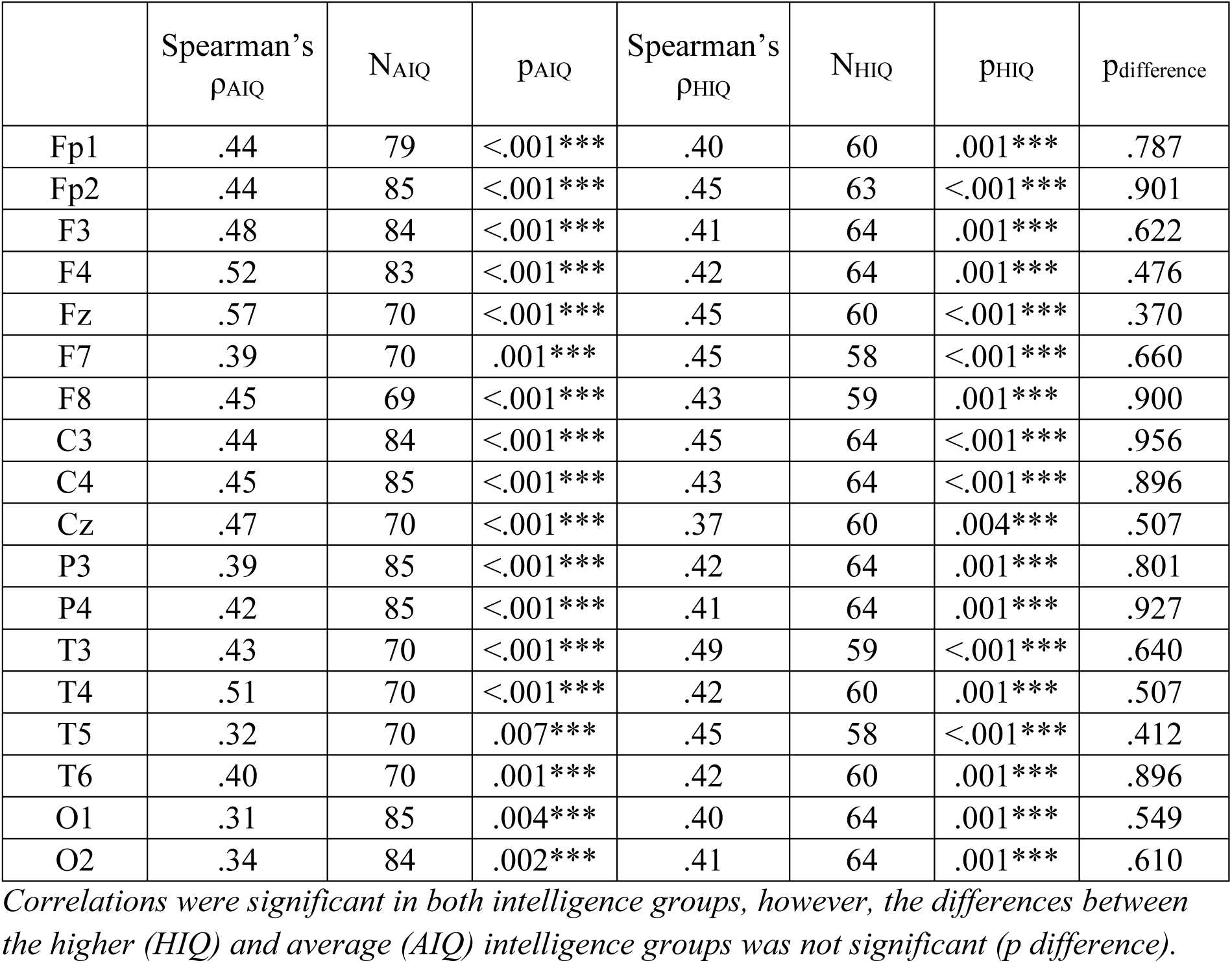
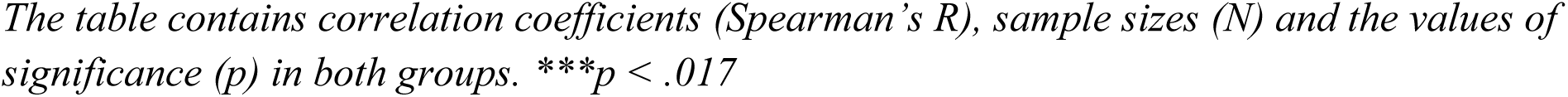
Comparison of the correlations between age and the slope of the NREM sleep EEG spectrum in subjects with average and high intelligence (AIQ vs HIQ)

| | Spearman's<br>$\rho_{AIQ}$ | $N_{AIQ}$ | $p_{AIQ}$ | Spearman's<br>$\rho_{HIQ}$ | $N_{HIQ}$ | $p_{HIQ}$ | $p_{\text{difference}}$ |
| --- | --- | --- | --- | --- | --- | --- | --- |
| Fp1 | .44 | 79 | <.001*** | .40 | 60 | .001*** | .787 |
| Fp2 | .44 | 85 | <.001*** | .45 | 63 | <.001*** | .901 |
| F3 | .48 | 84 | <.001*** | .41 | 64 | .001*** | .622 |
| F4 | .52 | 83 | <.001*** | .42 | 64 | .001*** | .476 |
| Fz | .57 | 70 | <.001*** | .45 | 60 | <.001*** | .370 |
| F7 | .39 | 70 | .001*** | .45 | 58 | <.001*** | .660 |
| F8 | .45 | 69 | <.001*** | .43 | 59 | .001*** | .900 |
| C3 | .44 | 84 | <.001*** | .45 | 64 | <.001*** | .956 |
| C4 | .45 | 85 | <.001*** | .43 | 64 | <.001*** | .896 |
| Cz | .47 | 70 | <.001*** | .37 | 60 | .004*** | .507 |
| P3 | .39 | 85 | <.001*** | .42 | 64 | .001*** | .801 |
| P4 | .42 | 85 | <.001*** | .41 | 64 | .001*** | .927 |
| T3 | .43 | 70 | <.001*** | .49 | 59 | <.001*** | .640 |
| T4 | .51 | 70 | <.001*** | .42 | 60 | .001*** | .507 |
| T5 | .32 | 70 | .007*** | .45 | 58 | <.001*** | .412 |
| T6 | .40 | 70 | .001*** | .42 | 60 | .001*** | .896 |
| O1 | .31 | 85 | .004*** | .40 | 64 | .001*** | .549 |
| O2 | .34 | 84 | .002*** | .41 | 64 | .001*** | .610 |

*Correlations were significant in both intelligence groups, however, the differences between the higher (HIQ) and average (AIQ) intelligence groups was not significant ( $p_{\text{difference}}$ ).*
*The table contains correlation coefficients (Spearman's $R$ ), sample sizes ( $N$ ) and the values of significance ( $p$ ) in both groups. \*\*\* $p < .017$*

### Overcoming model redundancy by determining the alternative intercept of the spectra

Although our model resulted in good fit with empirical data in terms of random (non-oscillatory) activity or coloured noise and the majority of our hypotheses (including the ones regarding peak power features) were supported by parameters derived from equation (2), the spectral slope and the intercept are far from being independent in statistical terms. That is, although women vs men differences emerged in our spectral intercepts (ln *C*_♀_ > ln *C*_♂_) as predicted in H4 (see Table 2), and no sex differences in NREM sleep EEG spectral slopes (*α*) were observed (Table 6a), the intercepts and the slopes are negatively correlated in our database (Table 6b): subjects with steeper spectral slopes (lower *α* exponents) are characterized by higher intercepts (apparently higher EEG amplitudes). This might reflect the position of the intercept, which is at ln *f* = 0 (*f* = 1 Hz). The interpolated 1 Hz power (based on the fitted line in the double logarithmic plots) partially reflects the steepness of the slope of the spectrum.

**Table 6.** Data on the lack of sex differences in spectral slopes and the relationship between spectral slopes and intercepts

| | a, sex differences in $\alpha$ | | | | | | b, $\alpha$ vs $\ln C_0$ | | |
| --- | --- | --- | --- | --- | --- | --- | --- | --- | --- |
| | t | p | N<sub>♀</sub> | $\overline{\alpha_{\text{♀}}}$ | N<sub>♂</sub> | $\overline{\alpha_{\text{♂}}}$ | r | p | N |
| Fp1 | 1.452 | .148 | 77 | -2.613 | 86 | -2.563 | -.83 | <.001*** | 163 |
| Fp2 | 1.477 | .141 | 81 | -2.624 | 90 | -2.575 | -.84 | <.001*** | 171 |
| F3 | 1.578 | .116 | 81 | -2.682 | 93 | -2.629 | -.83 | <.001*** | 174 |
| F4 | 1.585 | .114 | 81 | -2.689 | 92 | -2.634 | -.85 | <.001*** | 173 |
| Fz | 1.308 | .192 | 69 | -2.760 | 87 | -2.713 | -.85 | <.001*** | 156 |
| F7 | 1.307 | .192 | 67 | -2.510 | 86 | -2.460 | -.85 | <.001*** | 153 |
| F8 | 1.276 | .203 | 69 | -2.512 | 85 | -2.465 | -.83 | <.001*** | 154 |
| C3 | 2.044 | .042* | 81 | -2.658 | 93 | -2.594 | -.78 | <.001*** | 174 |
| C4 | 1.558 | .120 | 81 | -2.664 | 94 | -2.612 | -.81 | <.001*** | 175 |
| Cz | .878 | .381 | 69 | -2.701 | 87 | -2.672 | -.81 | <.001*** | 156 |
| P3 | 1.407 | .161 | 81 | -2.558 | 94 | -2.516 | -.79 | <.001*** | 175 |
| P4 | 1.398 | .163 | 81 | -2.553 | 94 | -2.510 | -.80 | <.001*** | 175 |
| T3 | 1.112 | .267 | 67 | -2.404 | 87 | -2.362 | -.80 | <.001*** | 154 |
| T4 | 1.031 | .303 | 69 | -2.400 | 87 | -2.361 | -.81 | <.001*** | 156 |
| T5 | .951 | .343 | 68 | -2.352 | 86 | -2.317 | -.80 | <.001*** | 154 |
| T6 | 1.358 | .176 | 68 | -2.359 | 87 | -2.312 | -.76 | <.001*** | 155 |
| O1 | 1.991 | .048* | 81 | -2.425 | 94 | -2.362 | -.79 | <.001*** | 175 |
| O2 | 2.038 | .043* | 81 | -2.430 | 93 | -2.369 | -.77 | <.001*** | 174 |

a. Sex differences in NREM sleep EEG spectral slopes as revealed by independent sample *t*-tests. The 3 descriptive significances do not survive the control of Type I error. b. The correlation of NREM sleep EEG spectral slopes ( $\alpha$ ) and intercepts at $\ln f = 0$ ( $\ln C_0$ ). Correlations are significant over the whole registered area and survive the control of multiple testing. Fisher *z*-transformed, averaged and back-transformed correlation value is -0.811.

In order to overcome the above issue of parameter-interdependency, we derived alternative intercepts with the aim of determining parts of the interpolated coloured spectrum at which our parameter do not reflects the steepness of the slope (*α*). We based our search for this alternative intercept on two assumptions: (1) the alternative (“slope-free”) intercept is situated at the border of low and high frequency activities, delineated by the reported sleep deprivation-induced increases and decreases of spectral power, respectively; (2) intercepts below the border mentioned in point 1. correlate negatively with the spectral slopes, whereas intercepts above this border correlate positively with slopes. Extended wakefulness of human adults is known to increase the NREM sleep EEG spectral power below the sleep spindle frequencies, that is the power of 1–9, 1–12 or 1–13 Hz according to different studies (Borbély et al., 1981; Finelli et al., 2001; Tinguely et al., 2006; Olbrich et al., 2014; Tarokh et al., 2015), whereas power above 10 or 13 Hz was shown to be decreased during recovery sleep (Finelli et al., 2001; Tinguely et al., 2006; Tarokh et al., 2015). Thus, we used our fitted model parameters *α* and ln *C* to determine the interpolated coloured power at frequencies of 7.4, 10, 12.2, 13.5, 15 and 20 Hz corresponding to natural logarithm values of 2, 2.3, 2.5, 2.6, 2.7, and 3, respectively. These alternative intercepts were tested for their independence from the slopes (*α*) by Pearson correlations (Fig 4). The pattern of correlations supported our assumptions: alternative intercepts below 12.2 Hz were found to correlate negatively with spectral slopes, whereas above 12.2 or 13.5 Hz (depending on electrode location) positive correlations were found. That is the best “slope-free” intercepts in the coloured part of the parametrized NREM sleep EEG spectra are emerging at 12.2 Hz and the 13.5 Hz for anterior and posterior derivations, respectively (ln *C*_*2*.*5*_ and ln *C*_*2*.*6*_). The original intercept derived at ln *f* = 0 could be termed as ln *C*_*0*_, according to this terminology. We reanalyzed H4 in terms of ln *C*_*2*.*5*_ and ln *C*_*2*.*6*_. The analyses resulted in increased mean effects sizes from 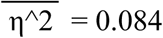 to 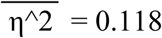 (both averaged over recording locations).

**Fig 4.**
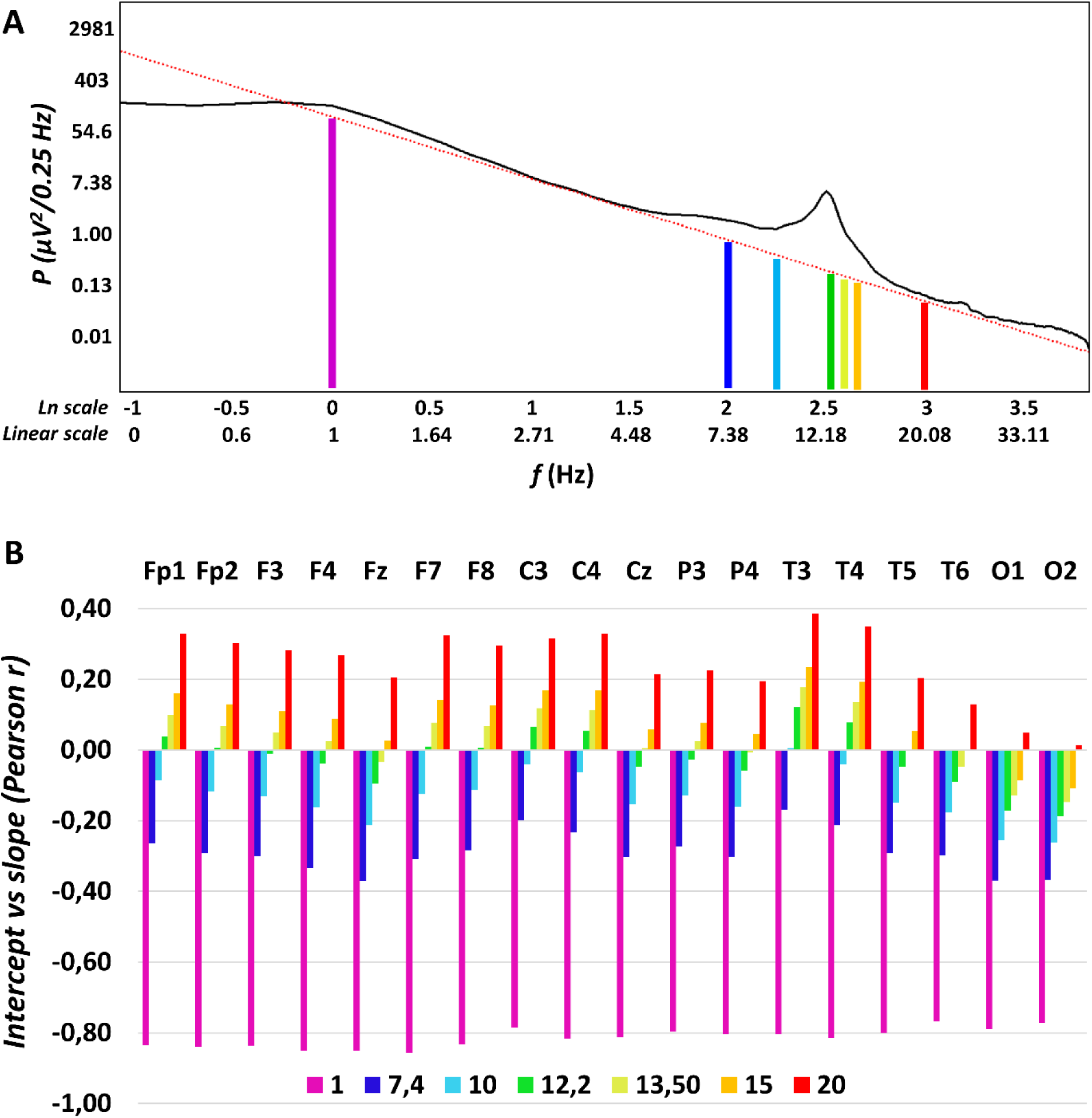
Determining the optimal alternative intercept for the NREM sleep EEG spectra. A. Linear fitted to the double logarithmic plot of an average NREM sleep EEG spectral power (P) derived from right frontopolar location (Fp2) in a young female volunteer. Beside the original, violet-colored intercept at ln f = 0 (f = 1 Hz), alternative intercepts are depicted at 7.4, 10, 12.2, 13.5, 15 and 20 Hz. B. Between-subject correlations of the potential intercepts (ln C) with the slopes of the spectra (α) in a location-dependent manner. Note the negative correlations for low and the positive correlation for high frequencies, respectively. Zero-correlations are seen in the middle of the sleep spindle frequency range (at 12.2 and 13.5 Hz), although occipital derivations are characterized by a slightly different pattern.

## Discussion

When analyzing the Fourier spectra of EEG records performed for long periods of sleep, researchers and clinicians rely on statistics. That is, the periodograms of short modified EEG segments are averaged in order to obtain the averaged spectra (Welch, 1967). As a consequence, the spectral profiles are inherently statistical in nature. The set of measures building up this statistical product conform to the power law functions characterized by negative exponents (Pereda et al., 1998; Freeman and Zhai, 2009), mixed up with a few positive deflections corresponding to non-random, oscillatory activity patterns (Lázár et al., 2020). In our view, the characterization of the Fourier spectrum by taking into account its electrophysiological and statistical regularities might result in an integrated characterization of NREM sleep EEG, which is superior in terms of construct validity and accuracy. First of all, a frequency-independent amplitude measure potentially reflecting non-neuronal factors, like skull anatomy, can be reliably separated and is not mixed up in power spectral values focusing on specific oscillatory phenomena. Although the natural logarithm of term C derived from formula (1) and (2) (ln *C*_*0*_) reliably reflects the hypothesized sex differences, the model could be refined by using alternative intercepts, which were independent from the slopes (ln *C*_*2*.*5*_ and ln *C*_*2*.*6*_) (Fig 4). The latter might constitute an ideal normalization value for NREM sleep EEG (spectra) in future basic and clinical studies.

In addition to the spectral intercepts, the power law functions describing the sleep EEG spectra appropriately address the issue of the ratio of EEG power at different frequencies, providing a single measure (*α*), instead of several ones scattered redundantly in all frequency bins and bands.

Last, but not least, spectral peak amplitudes (*P*_*Peak*_*(f)*) are whitened in our approach, that is, the coloured part of the spectrum is effectively controlled, which might enable researchers to differentiate random and non-random/oscillatory activities at specific frequencies.

The findings derived from our approach of parametrizing the NREM sleep EEG spectra clearly supports the robustness and validity of the method presented in this paper, which was inspired by studies aiming to whiten the spectral power in the sleep spindle frequency (Gottselig et al., 2002; Geiger et al., 2011). As predicted (H1), age correlates positively with NREM sleep EEG spectral exponents (Table 1a), indicating that aging is associated with less steep exponential decay slopes of the Fourier spectra (i.e. less negative exponents) (Fig 2A). This finding coheres with reports of bandwise power spectral analyses of NREM sleep EEG, indicating decreased low and increased high frequency activity in the NREM sleep EEG of healthy aged subjects (Carrier et al., 2001). Moreover, the steepness of the slope of the linear describing the relationship between the log-amplitude and the log-frequency of NREM sleep EEG revealed the same age-dependency (Feinberg et al., 1984). Thus, our method is capable of extracting spectral slope information with sufficient precision and is a valid and simple approach to be used in future (translational) studies. The slope of the spectrum is basically a measure of the constant ratio between low and high frequency activities, which was hypothesized to reflect the ratio between inhibition and excitation, the depth of sleep and/or the level of conscious awareness (Weiss et al., 2011; Gao et al., 2017; Colombo et al., 2019). Findings might indicate that aged subjects have lower sleep depth, but might also open new avenues beyond the exclusive focus on sleep slow waves/oscillation when studying the relationship between aging and sleep. The latter point is supported by our finding on the lack of a difference in the age-dependency of the NREM sleep EEG spectral slopes in subjects with average and high intelligence (Table 5). This finding apparently contrasts the outcomes of our previous report on the significant differences in age-dependent declines in NREM sleep EEG slow wave/oscillation of average and high IQ subjects. That is in terms of NREM sleep EEG slow waves high IQ subjects tend to age at a slower pace than average IQ subjects (Pótári et al., 2017). In spite of the fact that the database we used in the two studies are the same, the methods (classical spectral analysis vs. spectral exponent extraction) yield different results. That is, our present findings indicate that average and high IQ subjects tend to age at a same pace, at least in terms of their NREM sleep EEG spectral exponents. These contrasting results indicate that our former findings are preferentially reflecting the age- and IQ-dependency of the NREM sleep EEG slow oscillatory mechanism per se, but not the random activity and/or the constant ratio of slow and high frequency activities. The latter could be a subject of aging which is at least partially independent from the well characterized age-dependent decreases in slow oscillations (Mander et al., 2013) and is equally present in average and high IQ subjects. Recent findings and considerations suggest that the spectral slope derived from an electrophysiological signal indicates the ratio of excitation and inhibition in the underlying neural tissue (Gao et al., 2017). Thus, according to our current findings and previously published modeling data (Gao et al., 2017) aging is characterized by a relative increase in excitation over inhibition during the state of night time NREM sleep, and this effect seems to be relatively independent from the decreased slow oscillation reported in former studies (Mander et al., 2013; Pótári et al., 2017).

Aging was also shown to be associated with decreased sleep spindle frequency activity and decreased phasic sleep spindles in former studies (Purcell et al., 2017). These findings cohere with our current report of an age-associated decrease in whitened spectral peak amplitudes of NREM sleep EEG spindle frequency range (Table 1b). Reports suggest that the age-dependent decrease in sleep spindles recorded over the prefrontal regions mediates the cognitive decline in later ages (Mander et al., 2014). Moreover, it was suggested that this effect reflects the disruption of thalamocortical regulatory mechanisms involved in sleep spindle rhythmogenesis (Clawson et al., 2017). Thus, our index of whitened NREM sleep EEG spectral peak amplitude in the spindle frequency range could serve as a simple biomarker of the neurocognitive aspects of aging.

The age-associated increases in the frequency of sleep spindle oscillations (also known as intraspindle frequencies) were reported in several former reports (e.g. Principe and Smith, 1982), although the largest study did not reveal such changes in adulthood (Purcell et al., 2017). Our present findings reveal a non-predicted decrease in maximal frontal spectral peak amplitude in the spindle frequency range of NREM sleep EEG. The range of the spindle frequency changes clearly indicate a change from the predominant fast (∼14 Hz) to predominant slow (∼12 Hz) sleep spindle spectral peaks during aging. That is, our finding indicates a decrease in relative frontal emergence of fast sleep spindles during aging, rather than a deceleration of sleep spindles at a rate of 0.5 Hz/decade (Fig 2D). That is, our minimalistic goals to capture sleep spindle oscillatory activity with just one maximal spectral peak instead of two, resulted in the unexpected deceleration of sleep spindle frequency during aging in adult subjects.

Women were shown to be characterized by significantly higher NREM sleep EEG spectral intercepts as compared to men. This difference is not seen in the spectral slopes and is sharpened when using the alternative (“slope-free”) intercepts (ln *C*_*2*.*5*_ and ln *C*_*2*.*6*_ instead of ln *C*_*0*_). To the best of our knowledge this is the first report explicitly targeting these issues. We based our hypothesis on findings suggesting that women vs men differences in EEG power are largely frequency-independent (Carrier et al., 2001), thus indicating an overall amplitude effect captured by the term *C* in formula (1) and (2). That is, previous reports focusing on specific frequency ranges and oscillatory phenomena are confounded by overall amplitude differences in the EEG of women and men. Examples for such potentially confounded findings are reports on women vs men differences in sleep spindle densities/occurrences. Spindles detected by fixed thresholds (Crowley et al., 2002; Huupponene et al., 2002) or raw (non-whitened) spectral power values of the spindle frequency range (Dijk et al., 1989; Carrier et al., 2001) indicate sex differences (increased sleep spindle density/activity in women), but are not controlled for overall amplitude differences. It has to be noted however, that one of the early publications cited above hypothesized that women vs men differences in sleep EEG spectral power might reflect sex differences in skull thickness (Dijk et al., 1989), but - at least to our best knowledge - this hypothesis remained largely unexplored from the electrophysiological point of view. Our current approach considers this issue and provides a reliable and potentially useful method for controlling non-specific, non-neuronal effects in EEG amplitude. The estimation of the spectral intercept provides a simple index in the study of the skull-thickness-EEG power issue in future biophysical, electrophysiological-modeling studies. Our current findings clearly indicate the lack of sex differences in sleep spindle power when overall amplitude women vs men differences are controlled (Table 2).

Women were shown to be characterized by higher frequency sleep spindle oscillations as compared to men according to our former study based on the individual adjustment of sleep spindle frequencies and amplitudes (Ujma et al., 2014). This finding was strengthened by our current report based on the detection of whitened spectral peak location with .0052 Hz resolution (Table 3). That is, our current finding strengthens the validity of our spectral parametrization approach. In addition, the hypotheses suggesting that sleep spindle frequency is accelerated by either progesterone and its neuroactive, indirect GABA-agonist metabolite allopregnanolone (Driver et al., 1996) or the progesterone-induced hyperthermia (Deboer, 1998) during the follicular phase of the menstrual cycle in women are indirectly supported by our present findings. Although our participants were not controlled for menstrual cycle phases and oral contraceptive use, we can assume that at least some of the female subjects were examined during the follicular phase of their menstrual cycle. Furthermore, oral contraceptive use involve the intake of progestagenic compounds, which might induce some of the neural effects of endogenous progesterone in naturally cycling women.

Here we reveal a positive correlation between whitened spectral peak amplitude of sleep spindle frequency activity during NREM sleep and IQ in women, but not in men (Table 4; Fig 3). Intelligence was shown to be reflected in the intensity (amplitude and/or density) of phasic sleep spindle events or alternatively in the spectral power of sleep spindle frequency activity during NREM sleep (Bódizs et al., 2005; Ujma et al., 2014, Ujma et al., 2017; Ujma, 2018). In the database we use in our present study a marked sexual dimorphism of this effect was also revealed: women but not men were shown to be characterized by the sleep spindle amplitude/power vs IQ correlations (Ujma et al., 2014; Ujma et al., 2017). Although this latter effect was not unequivocally reflected in a significant meta-regression between effect size and % female in the sample in a subsequent metaanalysis, here we refer to it because convergent findings obtained by different methods used on the same dataset are an issue of validity of the methods. That is, we reproduced the positive sleep spindle vs. IQ correlation in women by using a linear fitting approach to the log-log spectra of NREM sleep and a concurrent whitening of spectral peaks, without assumptions on time-domain sleep spindle features. Again, this finding might strengthen our views on the reliability of the method of analyzing the constant, the slope and the (whitened) peak attributes of the NREM sleep EEG in human subjects.

Among the shortcomings of our work we would emphasize the lack of slow vs fast sleep spindle differentiation by the current version of our method, as well as the fact that we disregarded low frequency power (< 2 Hz) when fitting the slopes. Fitting of two slightly overlapping spectral peaks instead of just one, would increase considerably the complexity of the approach, whereas our intention was to keep the process as simple and intuitive as possible. Moreover, we intended to follow the already published method of finding the maximal peak in the spindle frequency range and correlating its amplitude/power with neurological-clinical and cognitive data (Gottselig et al., 2002; Geiger et al., 2011). Similarly, the potential and largely unpredictable contamination of low frequency power with sweating artefacts, as well as the high-pass filtering effects of gold-coated electrodes (Vanhatalo et al., 2005) we used in our studies precluded us from a precise measurement of the power law scaling at low frequencies below 2 Hz.

In sum, the parametrization of NREM sleep EEG of healthy adult subjects by relying on the power law scaling behavior of the electrical activity of the brain, as well as by completing this statistical property with the prominent spectral peak at the sleep spindle range, provides an integral method of describing and characterizing individual differences in sleep and cognition. Here we show, that most of the features of NREM sleep EEG can be efficiently compressed in the spectral intercepts, slopes and peaks, at least in terms of demographic (age, sex) and cognitive (IQ) correlates of sleep. It remains to be determined, if state-dependent changes, like overnight sleep dynamics and or sleep regulatory mechanisms can be appropriately described by these integrative parameters of NREM sleep. In addition, further studies are needed for an adequate handling of multiple spectral peaks and low frequency (< 2 Hz) oscillations in the non-full-band EEG.

## Methods

### Subjects/databases

Data was combined from multiple databases (Max Planck Institute of Psychiatry, Munich, Germany; Institute of Behavioural Sciences of Semmelweis University, Budapest, Hungary) for this retrospective multicenter study (Ujma et al., 2017; Ujma et al., 2019). Polysomnography data were recorded from 175 participants 81 females, 94 males, mean age 29.57 years, age range 17–60 years) and IQ scores were measured for 149 participants (68 females, 81 males, mean age 29.23 years, age range 17–60 years). Volunteers were recruited also via Mensa Germany and Mensa Hungary to increase the number of highly intelligent individuals. As some of the participants have missing data of some electrodes and/or IQ scores the data numbers from which the statistical analysis was conducted are always reported in the results.

Based on self-reports, none of the participants had a history of psychiatric or neurological disorders. Alcohol consumption was restricted before recording, but a small amount of caffeine (max. 2 cups of coffee before noon) was allowed to the participants. Based on self reports 8 participants were light or moderate smokers. Data were combined from multiple databases (Max Planck Institute of Psychiatry, Munich, Germany; Institute of Behavioural Sciences of Semmelweis University, Budapest, Hungary). The experiment was conducted in full accordance with the World Medical Association Helsinki Declaration and all applicable national laws and it was approved by the institutional review board, the Ethical Committee of the Semmelweis University, Budapest, or the Ludwig Maximilian University, Munich.

### Psychometric intelligence

Standardized nonverbal intelligence tests were recorded from 149 participants: 70 of them completed the Culture Fair Test (CFT) and 39 of them completed the Raven Advanced Progressive Matrices (Raven APM) test. 40 participants completed both tests. These tests have been shown to similarly measure abstract pattern completion and are particularly good measures of the general factor of intelligence. A composite raw intelligence test score was calculated, expressed as a Raven equivalent score (RES). RES for Raven APM tests was equal to the actual raw test score, whereas RES of the CFT test raw scores were equal to the Raven APM score corresponding to the IQ percentile derived from CFT performance and the age of the participant. Scores were averaged for participants who completed both tests. Standardization of APM was applied according to 1993 Des Moines (Iowa). Based on their mean IQ score, the sample was split into an average (AIQ: 88 < IQ < 120; 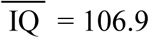; N = 85) and a high intelligence (HIQ: 120 ≤ IQ < 156; 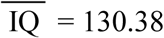; N = 64) subgroup (see Pótári et al., 2017).

### Polysomnography recordings

Detailed data recording procedures and power spectral analysis are also reported in the study of Ujma et al. (2019). Sleep data were recorded on two consecutive nights by standard polysomnography including EEG, electro-occulography (EOG), electrocardiography (ECG) and bipolar submental electromyography (EMG). EEG channels were placed according to the international 10–20 system (Fp1, Fp2, F3, F4, Fz, F7, F8, C3, C4, Cz, P3, P4, T3, T4, T5, T6, O1, O2 and left and right mastoids). Impedances for the EEG electrodes were kept below 8 kΩ. The sampling frequency was either 249 Hz, 250 Hz or 1024 Hz, depending on recording site (Supplementary table 1). All recordings were referenced to the mathematically linked mastoids. Data were offline re-referenced to the average of the mastoid signals and notch filtered at 50 Hz. Electrodes excluded from the analysis due to artifacts and/or recording failures were treated as missing data. The number missing data for the total 175 participants is reported in Supplementary Table 2, separately for each electrode. Recordings of the first night were used for habituation and therefore were not included in further analyses. Sleep data of the second night in the laboratory were scored for sleep-waking states and stages according to standard AASM criteria on a 20-sec basis (Iber et al., 2007) by an expert. Furthermore, artefactual segments were marked on a 4-sec basis and excluded from further analyses.

### Power spectral analysis

Power spectral densities were calculated for the NREM (N2 and N3) sleep, in .25 Hz bins from 0 Hz to the Nyquist frequency (*f*_*Nyquist*_) by relying on 4 s Hanning-tapered, non-artefactual windows. A 50% overlap was used for consecutive windows, whereas mixed-radix FFT calculating power spectral densities. Power spectral densities from all 4 s windows were then averaged. As data were recorded with different EEG devices producing different analog filter characteristics, average power spectral densities were corrected as follows: An analog waveform generator was connected to the C3 and C4 electrode positions of all EEG devices and sinusoid signals of various frequencies (0.05 Hz, every 0.1 Hz between 0.1–2 Hz, every 1 Hz between 2–20 Hz, every 10 Hz between 10–100 Hz) were generated with 40 and 355 μV amplitudes. The amplitude reduction rate of each recording system at each frequency was determined by calculating the proportion between digital (measured) and analog (generated) amplitudes of sinusoid signals at the corresponding frequency. The amplitude reduction rate was averaged for the 40 and 355 μV at each frequency. The reduction rate at the intermediate frequencies were interpolated by spline interpolation. The measured power spectral density values were corrected with the device-specific amplitude reduction rate by dividing the original value with the squared amplitude reduction rate at the corresponding frequency according to previous suggestions (Achermann and Borbély, 1997; Vasko et al., 1997).

### Estimation of the spectral intercepts and slopes

The power law function (formula (2)) was transformed to one which fits in the double logarithmic plots as follows (Fig. 1C):

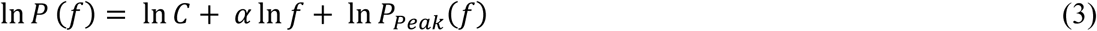

This means that the natural logarithm of spectral power (*P*) is expressed as a linear function of the natural logarithm of frequency (*f*). In addition, there are two terms in the equation: the natural logarithm of the constant (*C*) and the natural logarithm of peak power (*P*_*Peak*_, see Fig 1). If the latter equals 1 (*P*_*Peak*_ = 1), that is, there is no peak at a given frequency *f*, the value is 0 (ln 1 = 0). The logarithmic frequency scale inherently induces increasing data density at higher frequencies. Thus, a linear fit to this data would induce a strong bias against low frequency bins, which would contribute less to the determination of slopes compared to the higher frequency bins. In order to manage this problem and obtain an equal distribution of the data points, power values were interpolated up to the smallest frequency step (.0052 Hz) by the piecewise cubic Hermite interpolation method. In the next step a linear was fitted to the 2– 48 Hz frequency range of this equidistant log-log plot, excluding the 6.0052–17.9948 Hz frequency range corresponding to the alpha and spindle bands (in order to avoid those parts of the NREM sleep EEG spectra which are characterized by non-random, oscillatory activities as well). This part of our procedure was inspired by two former studies using a similar approach for whitening of the sleep spindle spectra (Gottselig et al., 2002; Geiger et al., 2011). The slope of the linear is *α*, whereas its intercept is ln *C*.

### Estimation of the spectral peak frequencies

Spectral peak frequency was determined in the 9–18 Hz range, separately for each EEG derivation by automatically defining local maxima in mathematical terms. That is, we used the first derivative test in order to find the critical points, followed by the second derivative test to differentiate local maxima and minima. A spectral peak was accepted if the first order derivative was zero and the second order derivative was negative. Calculations were performed as follows: a second-degree polynomial curve fitting was performed using all sets of successive bin triplets (.75 Hz), with an overlap of 2 bins (.5 Hz) in the 9–18 Hz range resulting in equations of the following type:

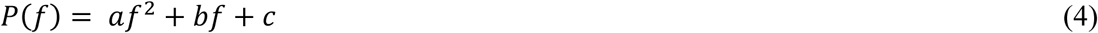

*P*: power

*f*: frequency (9–18 Hz)

*a, b*, and *c*: fitted parameters.

The first derivative of these functions were calculated for each triplet, resulting in:

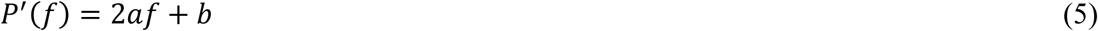

The slope of the function described in formula (5) is *2a*, which was considered as the derivative at the middle of the triplets, resulting in the first derivative function of the spectra. The procedure was repeated for calculating the second derivatives: in this case the first order derivative function served as an input for fitting the quadratic polynomials.

Zero-crossings of the first derivatives were determined by spline interpolation (interpolating the series between the bins of .25 Hz). In addition, the second derivative was interpolated by the spline method at each detected zero crossing of the first derivatives. The cases which were characterized by the co-ocurrences of the two criteria below were considered as spectral peak frequencies:

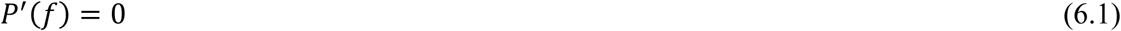

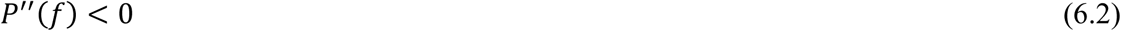

### Estimation of the spectral peak amplitudes

Spectral power at peak frequencies were estimated by spline interpolation of the double logarithmic plots of the power spectra. The spectral peak amplitude was then whitened by subtracting the estimated power based on the fitted linear function from the coloured peak power:

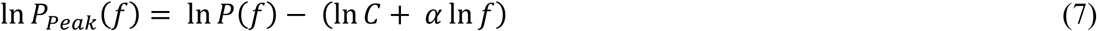

In order to avoid negative amplitudes due to the logarithmic scale, the power values were shifted for being all positive before this subtraction by adding a constant. This latter step was applied for the calculation of the amplitude measures only. As multiple spectral peaks were detected for some of the participants/EEG derivations, the one with the largest amplitude was determined and used in this study. If no spectral peak was found in the spindle frequency range, peak values were considered as missing data (see Table 5.). Data analysis was performed by Matlab R2018b (Mathworks Inc.).

### Statistical analyses

Goodness of fit of the linear to the equidistant log-log spectral data was assessed by Pearson product moment correlations, which were Fisher Z-transformed, averaged and back-transformed according to Silver and Dunlap (1987). Last, but not least the resulting average R-value were squared in order to determine the shared variance. Standard deviation (SD) was assessed from the Fisher-Z-transformed dataset, and the resulting value was back-transformed as well.

We used parametric tests (Pearson correlation, independent sample t-test) on normally distributed data and non-parametric tests (Spearman’s rank correlation, Mann-Whitney U test) when the distribution of the data was not Gaussian. The normality of the distributions was analysed by Shapiro-Wilk tests. In order to control Type 1 statistical errors due to multiple electrodes/hypothesis, we used a version of the Descriptive Data Analysis (DDA) protocol (Abt, 1987) adapted for neurophysiological data (Abt, 1990; Duffy, 1990). This procedure tests the global null hypothesis (“all individual null hypotheses in the respective region are true”) at level .05, against the alternative that at least one of the null hypotheses is wrong. DDA considers the intercorrelations between the different electrodes and is based on defining Rüger’s areas (Rüger, 1978), which are sets of spatially contingent conventionally (descriptively) significant (p < .05) results. The global significance of the Rüger area means that at least 1/3 of the descriptive significances are significant at a p = .05/3 = .017 and/or ½ of the descriptive significances are significant at p = .05/2 = .025. We used both criteria simultaneously (the “and” operator) in this study. In order to obtain a better localization of regions with significant correlations, associations between NREM sleep EEG spindle frequency whitened spectral peak amplitudes and IQ were represented by significant probability maps (Hassainia et al., 1994).

## Acknowledgements

We would like to thank Mensa Germany and Mensa Hungary for their help in volunteer recruitment.

## Supporting information

**Supplementary table 1.**
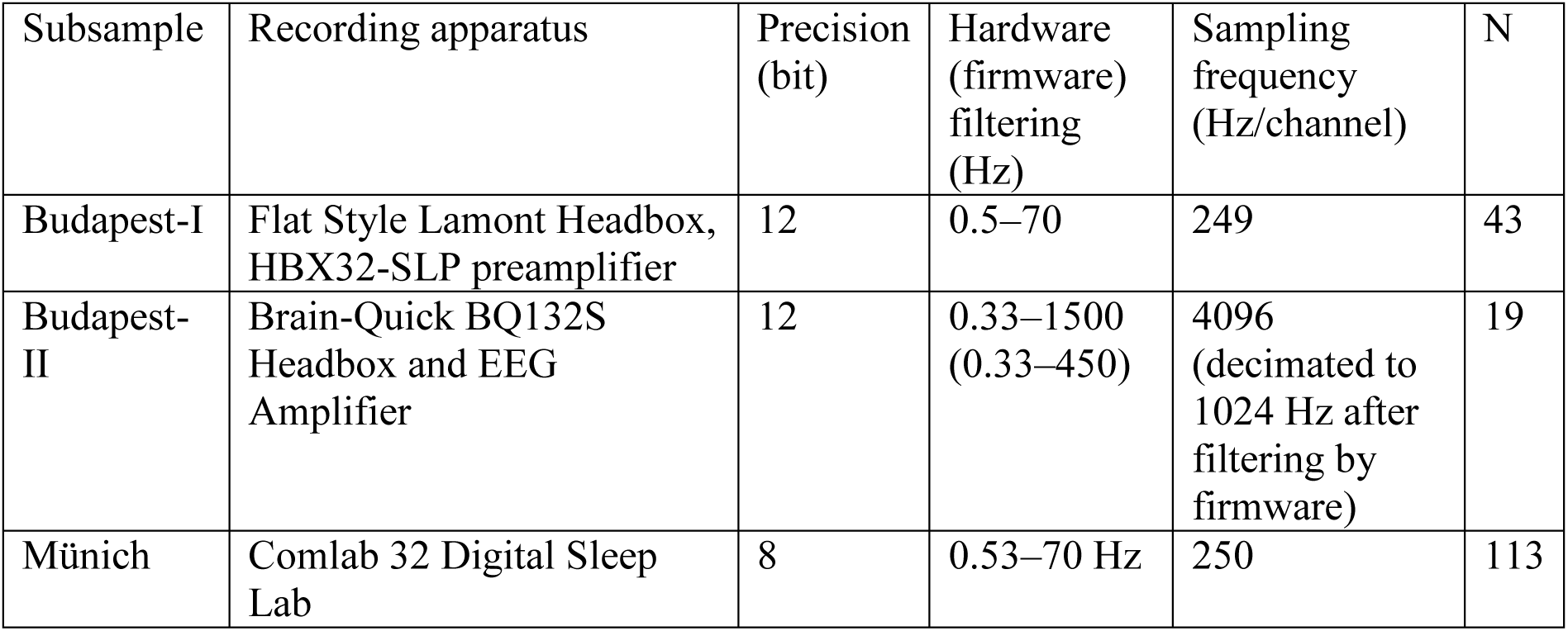
Technical details of the recordings in different subsamples included in the present investigation

**Supplementary table 2.**
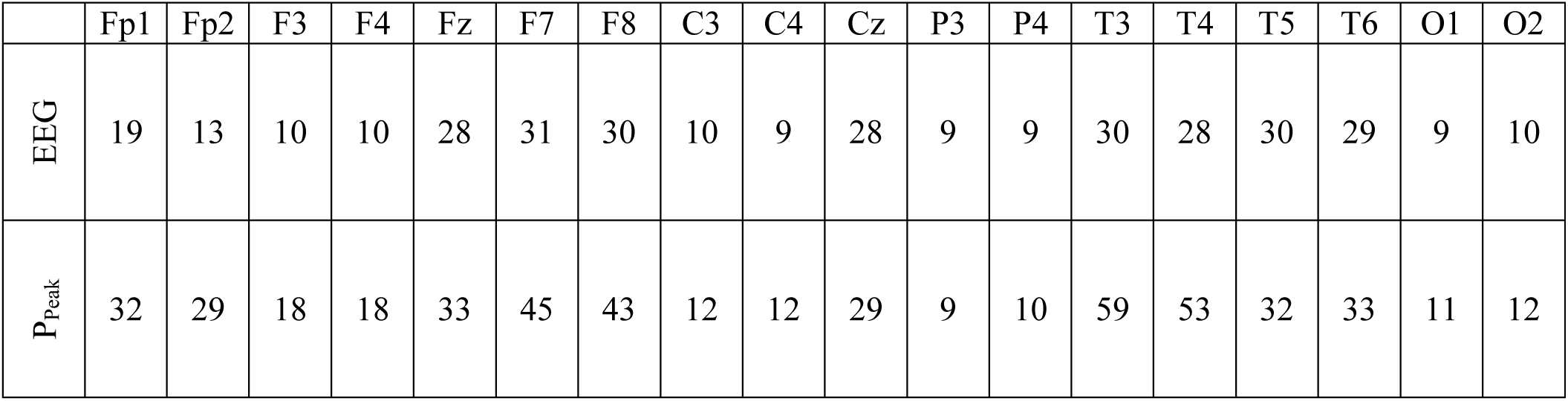
The number of missing/artefactual records (EEG) and peak power values (P_Peak_), separately for each electrode

## References

Abt K. Descriptive data analysis: a concept between confirmatory and exploratory data analysis. Methods Inf Med. 1987;26(02):77–88. doi: 10.1055/s-0038-1635488

Abt K. Statistical aspects of neurophysiologic topography. J Clin Neurophysiol. 1990;7(4):519–534. doi: 10.1097/00004691-199010000-00007

Achermann P, Borbély AA. Low-frequency (< 1 Hz) oscillations in the human sleep electroencephalogram. Neuroscience. 1997;81(1):213–222. doi: 10.1016/S0306-4522(97)00186-3

Bódizs R, Kis T, Lázár AS, Havrán L, Rigó P, Clemens Z, et al. Prediction of general mental ability based on neural oscillation measures of sleep. J Sleep Res. 2005;14(3):285–292. doi: 10.1111/j.1365-2869.2005.00472.x

Bódizs R, Gombos F, Kovács I. Sleep EEG fingerprints reveal accelerated thalamocortical oscillatory dynamics in Williams syndrome. Res Dev Disabil. 2012;33(1):153–164. doi: 10.1016/j.ridd.2011.09.004

Borbély AA, Baumann F, Brandeis D, Strauch I, Lehmann D. Sleep deprivation: effect on sleep stages and EEG power density in man. Electroencephalogr Clin Neurophysiol. 1981;51(5):483–495. doi: 10.1016/0013-4694(81)90225-x

Campbell IG, Grimm KJ, de Bie E, Feinberg I. Sex, puberty, and the timing of sleep EEG measured adolescent brain maturation. Proc Natl Acad Sci U S A. 2012;109(15):5740–5743. doi: 10.1073/pnas.1120860109

Carrier J, Land S, Buysse DJ, Kupfer DJ, Monk TH. The effects of age and gender on sleep EEG power spectral density in the middle years of life (ages 20-60 years old). Psychophysiology. 2001;38(2):232–242. doi: 10.1111/1469-8986.3820232

Clawson BC, Durkin J, Aton SJ. Form and function of sleep spindles across the lifespan. Neural Plast. 2016;2016:6936381. doi: 10.1155/2016/6936381

Colombo MA, Napolitani M, Boly M, Gosseries O, Casarotto S, Rosanova M, et al. The spectral exponent of the resting EEG indexes the presence of consciousness during unresponsiveness induced by propofol, xenon, and ketamine. Neuroimage. 2019;189:631–644. doi: 1.1016/j.neuroimage.2019.01.024.

Crowley K, Trinder J, Kim Y, Carrington M, Colrain IM. The effects of normal aging on sleep spindle and K-complex production. Clin Neurophysiol. 2002;113(10):1615–1622. doi: 10.1016/s1388-2457(02)00237-7

Deboer T. Brain temperature dependent changes in the electroencephalogram power spectrum of humans and animals. J Sleep Res. 1998;7(4):254–262. doi: 10.1046/j.1365-2869.1998.00125.x

Dijk DJ, Beersma DG, Bloem GM. Sex differences in the sleep EEG of young adults: visual scoring and spectral analysis. Sleep. 1989;12(6):500–507. doi: 10.1093/sleep/12.6.500

Driver HS, Dijk DJ, Werth E, Biedermann K, Borbély AA. Sleep and the sleep electroencephalogram across the menstrual cycle in young healthy women. J Clin Endocrinol Metab. 1996;81(2):728–735. doi: 10.1210/jcem.81.2.8636295

Duffy FH, Jones K, Bartels P, Albert M, McAnulty GB, Als H. Quantified neurophysiology with mapping: statistical inference, exploratory and confirmatory data analysis. Brain Topogr. 1990;3(1):3–12. doi: 10.1007/bf01128856

Feinberg I, March JD, Fein G, Aminoff MJ. Log amplitude is a linear function of log frequency in NREM sleep eeg of young and elderly normal subjects. Electroencephalogr Clin Neurophysiol. 1984;58(2):158–116. doi: 10.1016/0013-4694(84)90029-4

Freeman WJ, Holmes MD, West GA, Vanhatalo S. Fine spatiotemporal structure of phase in human intracranial EEG. Clin Neurophysiol 2006;117(6):1228–1243. doi: 10.1016/j.clinph.2006.03.012

Freeman WJ, Zhai J. Simulated power spectral density (PSD) of background electrocorticogram (ECoG). Cogn Neurodyn. 2009;3(1):97–103. doi: 1.1007/s11571-008-9064-y

Gao R, Peterson EJ, Voytek B. Inferring synaptic excitation/inhibition balance from field potentials. Neuroimage 2017;158:70–78. doi: 10.1016/j.neuroimage.2017.06.078

Geiger A, Huber R, Kurth S, Ringli M, Jenni OG, Achermann P. The sleep EEG as a marker of intellectual ability in school age children. Sleep. 2011;34(2):181–189. doi: 10.1093/sleep/34.2.181

Gottselig JM, Bassetti CL, Achermann P. Power and coherence of sleep spindle frequency activity following hemispheric stroke. Brain. 2002;125(Pt 2):373–383. doi: 10.1093/brain/awf021

Hassainia F, Petit D, Montplaisir J. Significance probability mapping: the final touch in t-statistic mapping. Brain Topogr. 1994;7(1):3–8. doi: 10.1007/bf01184832

Huupponen E, Himanen SL, Värri A, Hasan J, Lehtokangas M, Saarinen J. A study on gender and age differences in sleep spindles. Neuropsychobiology. 2002;45(2):99–105. doi: 10.1159/000048684

Iber C, Ancoli-Israel S, Chesson A, Quan SF, eds. The AASM manual for the scoring of sleep and associated events: rules, terminology, and technical specification, 1st ed. Westchester, IL: American Academy of Sleep Medicine, 2007.

Kaskie RE, Ferrarelli F. Sleep disturbances in schizophrenia: what we know, what still needs to be done. Curr Opin Psychol. 2019;34:68–71. doi: 1.1016/j.copsyc.2019.09.011.

Lázár AS, Lázár ZI, Bódizs R. Frequency Characteristics of Sleep. In: Oxford Handbook of EEG frequency. (in press)

Looker AC, Melton LJ 3rd, Harris T, Borrud L, Shepherd J, McGowan J. Age, gender, and race/ethnic differences in total body and subregional bone density. Osteoporos Int. 2009;20(7):1141–1149. doi: 10.1007/s00198-008-0809-6

Mander BA, Rao V, Lu B, Saletin JM, Lindquist JR, Ancoli-Israel S, et al. Prefrontal atrophy, disrupted NREM slow waves and impaired hippocampal-dependent memory in aging. Nat Neurosci. 2013;16(3):357–364. doi: 10.1038/nn.3324

Mander BA, Rao V, Lu B, Saletin JM, Ancoli-Israel S, Jagust WJ, et al. Impaired prefrontal sleep spindle regulation of hippocampal-dependent learning in older adults. Cereb Cortex 2014;24(12):3301–3309. doi: 10.1093/cercor/bht188

Nicolas A, Petit D, Rompré S, Montplaisir J. Sleep spindle characteristics in healthy subjects of different age groups. Clin Neurophysiol. 2001;112(3):521–527. doi: 10.1016/s1388-2457(00)00556-3

Olbrich E1, Landolt HP, Achermann P. Effect of prolonged wakefulness on electroencephalographic oscillatory activity during sleep. J Sleep Res. 2014;23(3):253–260. doi: 10.1111/jsr.12123.

Pereda E, Gamundi A, Rial R, Gonzalez J. Non-linear behaviour of human EEG: fractal exponent versus correlation dimension in awake and sleep stages. Neurosci Lett. 1998;250(2):91–94. doi: 10.1016/s0304-3940(98)00435-2

Pótári A, Ujma PP, Konrad BN, Genzel L, Simor P, Körmendi J, et al. Age-related changes in sleep EEG are attenuated in highly intelligent individuals. Neuroimage. 2017;146:554–560. doi: 1.1016/j.neuroimage.2016.09.039

Principe JC, Smith JR. Sleep spindle characteristics as a function of age. Sleep. 1982;5:73–84. doi: 10.1093/sleep/5.1.73

Purcell SM, Manoach DS, Demanuele C, Cade BE, Mariani S, Cox R, et al. Characterizing sleep spindles in 11,630 individuals from the National Sleep Research Resource. Nat Commun. 2017;8:1593. doi: 1.1038/ncomms1593.

Rüger B. Das maximale Signifikanziniveau des tests: “Lehne H0 ab, wenn k unter n gegebene Tests zur ablehnung führen”. Metrika 1978;25:171–178. doi: 10.1007/bf02204362

Silver NC, Dunlap WP. Averaging correlation coefficients: Should Fisher’s z transformation be used? J Appl Psychol. 1987;72(1):146–148. doi: 10.1037/0021-9010.72.1.146

Tarokh L, Rusterholz T, Achermann P, Van Dongen HP. The spectrum of the non-rapid eye movement sleep electroencephalogram following total sleep deprivation is trait-like. J Sleep Res. 2015;24(4):360–363. doi: 10.1111/jsr.12279.

Tinguely G, Finelli LA, Landolt HP, Borbély AA, Achermann P. Functional EEG topography in sleep and waking: state-dependent and state-independent features. Neuroimage. 2006;32(1):283–292.

Ujma PP, Konrad BN, Genzel L, Bleifuss A, Simor P, Pótári A, et al. Sleep spindles and intelligence: evidence for a sexual dimorphism. J Neurosci. 2014;34(49):16358–16368. doi: 1.1523/JNEUROSCI.1857-14.2014

Ujma PP, Konrad BN, Gombos F, Simor P, Pótári A, Genzel L, et al. The sleep EEG spectrum is a sexually dimorphic marker of general intelligence. Sci Rep. 2017;7(1):1807. doi: 10.1038/s41598-017-18124-0

Ujma PP. Sleep spindles and general cognitive ability–A meta-analysis. Sleep Spindles & Cortical Up States, 2018. doi: 10.1556/2053.2.2018.01

Ujma PP, Konrad BN, Simor P, Gombos F, Körmendi J, Steiger A, et al. Sleep EEG functional connectivity varies with age and sex, but not general intelligence. Neurobiol Aging. 2019;78:87–97. doi: 10.1016/j.neurobiolaging.2019.02.007

Ujma PP, Simor P, Steiger A, Dresler M, Bódizs R. Individual slow-wave morphology is a marker of aging. Neurobiol Aging. 2019;80:71–82. doi: 10.1016/j.neurobiolaging.2019.04.002

Ujma PP, Bódizs R, Dresler M. Sleep and intelligence: critical review and future directions. Curr Opin Behav Sci. 2020;33:109–117. doi: 10.1016/j.cobeha.2020.01.009

Vanhatalo S, Voipio J, Kaila K. Full-band EEG (FbEEG): an emerging standard in electroencephalography. Clin Neurophysiol. 2005;116(1):1–8. doi: 10.1016/j.clinph.2004.09.015

Vasko RC Jr, Brunner DP, Monahan JP, Doman J, Boston JR, el-Jaroudi A, et al. Power spectral analysis of EEG in a multiple-bedroom, multiple-polygraph sleep laboratory. Int J Med Inform. 1997;46(3):175–184. doi: 10.1016/s1386-5056(97)00064-6

Weiss B, Clemens Z, Bódizs R, Halász P. Comparison of fractal and power spectral EEG features: Effects of topography and sleep stages. Brain Res Bull. 2011;84(6):359–375. doi: 10.1016/j.brainresbull.2010.12.005

Welch PD. The use of Fast Fourier Transform for the estimation of power spectra: A method based on time averaging over short, modified periodograms. IEEE Transactions on Audio and Electroacoustics. 1967;15(2):70–73. doi: 10.1109/TAU.1967.1161901

